# Loose coupling between Ca^2+^ channels and release sensors as a synaptic correlate of higher order brain function

**DOI:** 10.64898/2025.12.15.694355

**Authors:** Grit Bornschein, Max Schwarze, Antonia Brunner, Akanksha Arshia, Simone Brachtendorf, Hartmut Schmidt

## Abstract

In the mature neocortex, functionally distinct areas are built by the same archetypes of neurons, but depending on the area, these neurons and their synapses are engaged in very different functions, ranging from lower order processing of sensory information to higher order associations and cognitive functions. We found significant differences in the functional presynaptic nanoarchitectures of the same types of pyramidal neuron synapses, depending on whether they are located in the prefrontal cortex (PFC) or in the primary somatosensory cortex (S1). Synapses in PFC operated with loose microdomain coupling as opposed to tight nanodomain coupling in S1. These differences were associated with significant differences in synaptic timing, efficacy and plasticity between areas. Our data suggest that the mature neocortex uses tuning of synaptic topographies to specialize seemingly identical types of neurons for their required function. They suggest that microdomain coupling in pyramidal neuron synapses could be a presynaptic structure-function correlate of higher order neocortical functions.

## Introduction

The neo- or isocortex is a morphologically homogeneous brain region that covers highly diverse functions within its specialized areas, ranging from early sensory processing and motor control up to higher order associations, cognition and consciousness (Douglas and Martin, 2004; Harris and Shepherd, 2015). Notably, all of these diverse functions are executed by the same archetypes of neurons and synapses. However, depending on the area, they are engaged to different degrees in lower or higher order processing. Whether area-specific functional differences in the mature neocortex are associated with or even arise from differences in the functional presynaptic nanoarchitecture of the same principal types of synapses is currently unclear.

At presynaptic active zones, the coupling distance between voltage-gated Ca^2+^ channels (VGCCs) and transmitter-filled synaptic vesicles (SVs) is a major determinant of key synaptic properties, including speed, efficacy and reliability of synaptic transmission (Rozov et al., 2001; Fedchyshyn and Wang, 2005; Bucurenciu et al., 2008; Baur et al., 2015; Bornschein et al., 2019b; Chen et al., 2024), and also short-term plasticity, although the latter can be obscured by vesicle replenishment (Miki et al., 2016; Doussau et al., 2017; Bornschein et al., 2019a; Lin et al., 2022). Studies at different synapses in different parts of the brain suggest that excitatory synapses engaged in reliable information transfer switch from loose microdomain to tight nanodomain coupling during development (Fedchyshyn and Wang, 2005; Baur et al., 2015; Nakamura et al., 2015; Bornschein et al., 2019b). This includes synapses between layer 5 pyramidal neurons (L5PNs) in the primary somatosensory cortex (S1), an area of early sensory processing (Bornschein et al., 2019b). On the other hand, loose microdomain coupling was found in the mature brain to date only at a highly plastic hippocampal synapse and has been suggested to provide a molecular framework for presynaptic plasticity (Vyleta and Jonas, 2014). However, if microdomain coupling also represents a synaptic correlate of higher order neocortical function or if it restricted to specific and highly plastic synapses in the hippocampus is currently unclear. Therefore, here we tested the hypothesis that loose microdomain coupling is the synaptic correlate of higher order functions of the mature neocortex, whereas tight nanodomain coupling is the synaptic correlate of earlier processing stages. Our data show that in the mature cortex the same types of pyramidal neuron synapses operate with loose microdomain coupling if they are located in the prefrontal cortex (PFC), whereas in S1 they operate with tight nanodomain coupling. Thus, our results corroborate the hypothesis and suggest microdomain coupling as a synaptic correlate of higher order function in the neocortex.

## Results

### Synaptic efficacy, reliability and short-term plasticity differ between PFC and S1

We focused on the lateral prefrontal cortex (PFC) and medial prefrontal cortex (mPFC) as areas of higher order processing and on primary somatosensory cortex (S1) as an area of lower order processing. Within these areas we investigated excitatory synapses between pyramidal neurons (PNs), which are the principal building blocks of the neocortex. We first analyzed the properties of synapses connecting neighboring layer 5 PNs (L5PNs) in mature PFC (P21-26) by performing paired whole-cell patch-clamp recordings and compared their properties to those we previously reported for the same synapses and age in S1 (Bornschein et al., 2019a; Bornschein et al., 2019b) (**Figure 1*A*, Table 1**). S1 data from both previous studies were combined and additional analyses of synaptic delays, PPRs and success rates were performed. We found that the efficacy of transmission, i.e. the size of a unitary EPSC evoked by a single action potential (Rozov et al., 2001), was significantly lower in PFC (8 pA, 5-15 pA) than in S1 (29 pA, 18-53 pA), whereas the synaptic failure rates were significantly higher in PFC (0.16, 0.08-0.28) than in S1 (0, 0-0.03; **Figure 1*B-D***). In addition, we found that the synaptic delays (2.46 ms, 2.20-2.94 ms) as well as their standard deviation (SD_Delay_: 0.60 ms, 0.49-0.81 ms) were significantly larger in PFC than in S1 (delay: 1.99 ms, 1.67-2.25 ms; SD_Delay_: 0.24 ms, 1.14-0.37 ms; **Figure 1*E, F***), indicating that transmitter release is less tightly coupled to the time of the action potential in PFC than in S1 (Bullmann et al., 2024). Together these data show that synaptic speed, efficacy and reliability during single action potentials are lower in PFC than at the same synapses in S1.

**Table 1.** New and previously published data.

| Synapse / region | Experiments / analyses | Figures | Data |
| --- | --- | --- | --- |
| L5PN-L5PN PFC | EPSC amplitudes, failure rates, delays, $SD_{Delay}$, PPRs, success rates | Figure 1B-I<br>Figure S1A,B | This study |
| L5PN-L5PN S1 | EPSC amplitudes, failure rates; | Figure 1B-I | Bornschein et al., |
| | additional quantification of delays, $SD_{\text{DelayS}}$, PPRs, success rates | **Figure S1B** | 2019a, b |
| L2/3-L5PN PFC and S1 | PPRs, $SD_{\text{Delay}}$ | **Figure 1J-L**<br>**Figure S1C,D** | This study |
| L2/3-L5PN PFC and S1 | NMDA receptor contribution | **Figure S2A-F** | This study |
| L5PN-L5PN PFC | Chelator experiments, CD | **Figure 2A,B** | This study |
| L5PN-L5PN S1 | Chelator experiments, CD;<br>Reanalyzes of EGTA effects at 20-30 min test period | **Figure 2B** | Bornschein et al., 2019b |
| L2/3-L5PN PFC and S1 | Chelator experiments, CD | **Figure 2C-F**<br>**Figure S3A-E** | This study |
| L5PN-L5PN PFC | MPFA | **Figure 3A,D,E** | This study |
| L5PN-L5PN S1 | MPFA | **Figure 3D,E** | Bornschein et al., 2019b |
| L2/3-L5PN PFC and S1 | MPFA | **Figure 3B,C,F** | This study |
| L5PN PFC | 2P $Ca^{2+}$-Imaging | **Figure 4A-F** | This study |
| L5PN S1 | 2P $Ca^{2+}$-Imaging | **Figure 4A-F**<br>**Figure S4A** | This study |
| L5PN S1 | 2P $Ca^{2+}$-Imaging: basal $[Ca^{2+}]_i$ with Fluo-5F and OGB-1 | **Figure S4B** | Bornschein et al., 2025; this study |
| L2/3PN and L5PN PFC | Modeling | **Figure 5A-D**<br>**Figure S5** | This study |
| L2/3-L5PN PFC | $Cd^{2+}$ experiments | **Figure 5E,F** | This study |
| L2/3-L5PN PFC | $Cd^{2+}$ experiments | **Figure 5E,F** | This study |

**Figure 1.**
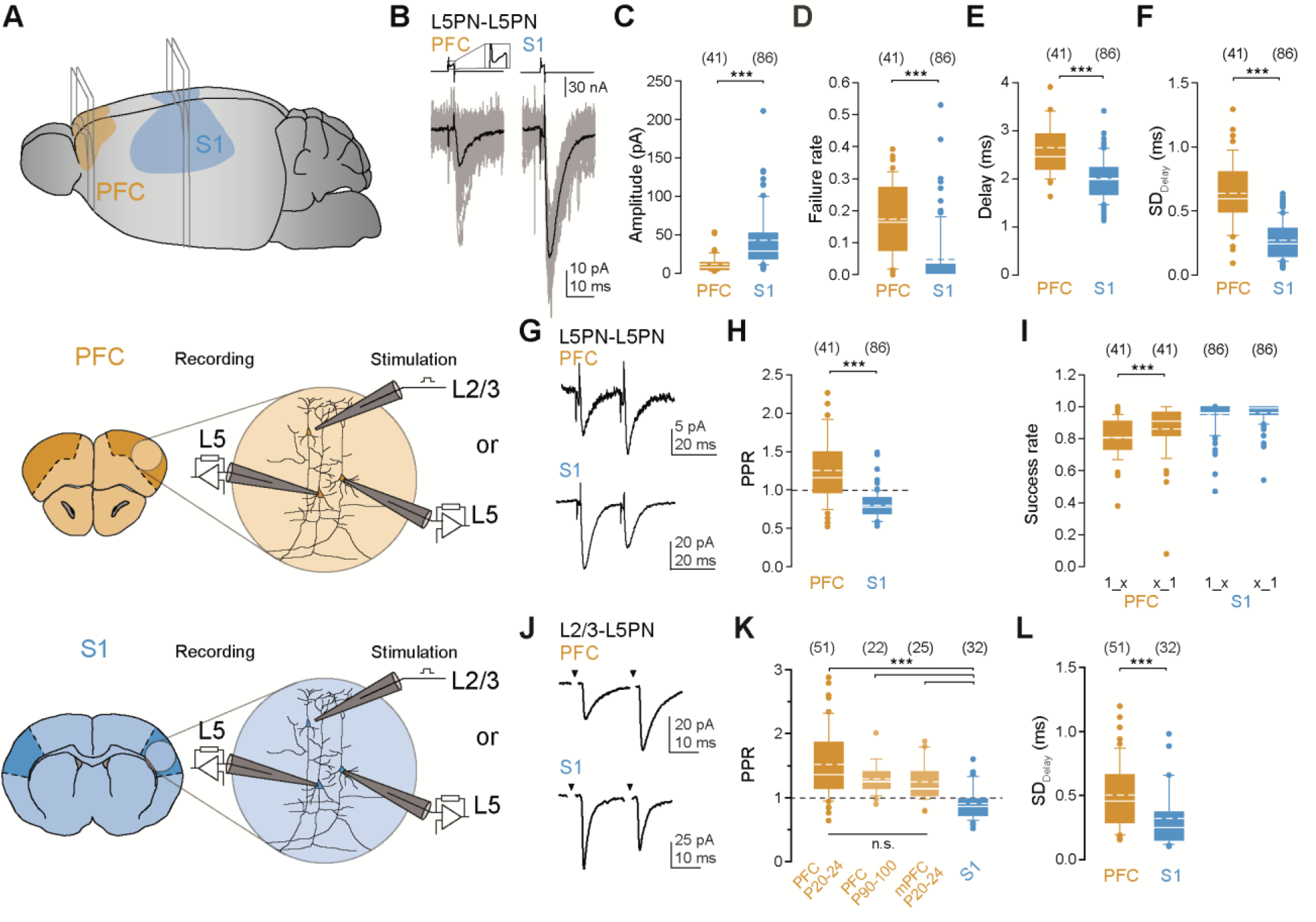
Differences in synaptic transmission and plasticity between PFC and S1 synapses. ***(A)***Schematic representation of the mature mouse brain (top) and the positions at which coronal sections were made from PFC (orange; middle) or S1 (blue; bottom). Recordings were made in acute slices from L5PN-L5PN pairs or from L2/3-L5PN connections. ***(B)***EPSCs (bottom) recorded from postsynaptic L5PNs after evoking action currents in the presynaptic L5PNs (top, inset: 3fold magnification) in PFC (left) or S1 (right), individual recordings in gray, averages in black). **(*C-F*)** Summary of basic synaptic properties: EPSC amplitudes (***C***), failure rates (***D***), synaptic delays (***E***) and their SD values (***F***) in PFC and S1 (medians ± IQRs, means as dashed lines, whiskers represent 10th and 90th percentiles, dots indicate outliers, numbers of cell pairs in brackets; ***P<0.001, Mann-Whitney U test (MWU)). Data from S1 are from ref. (Bornschein et al., 2019b). **(*G*)** EPSCs recorded from pairs of L5PNs at 50 Hz intervals in PFC (top) and S1 (bottom), presynaptic stimulations are omitted, averages from at least 10 individual recordings). **(*H, I*)** Summary of PPRs (**H**; ***P<0.001, MWU) and success rates (**I**) following the 1st (1_x) and 2nd stimulation (x_1; ***P<0.001, P=0.987, Wilcoxon signed rank test (WSR)). S1 data are from ref. (Bornschein et al., 2019a). **(*J*)** As in (***G***) but for L2/3-L5 synapses (bottom, averages from at least 15 individual recordings, stimulation artefacts were blanked for clarity, arrowheads denote the time points of extracellular stimulation). **(*K*)** Boxplots summarizing PPR data from lateral PFC (PFC), medial PFC (mPFC), older PFC (P90-100) and in S1 (***P<0.001, n.s., not significant, ANOVA on ranks). **(*L*)** Comparison of SDs of the synaptic delays at L2/3-L5 synapses between PFC and S1 (***P<0.001, MWU).

When pairs of action potentials were applied at high frequency (**Figure 1*G-I***), differences in the synaptic success rates were associated with a significant difference in the paired pulse ratios (PPR) between PFC and S1. In PFC paired pulse facilitation (PPF, 1.18, 0.98-1.52) prevailed associated with increased success rates in the 2^nd^ pulse (1_x: 0.81, 0.73-0.91; x_1: 0.91, 0.82-0.97). In contrast, in S1 paired pulse depression (PPD, 0.78, 0.69-0.90) was found with high initial success rates (1_x: 1.00, 0.97-1.00; x_1: 0.99, 0.95-1.00). We observed the difference in short-term plasticity over a range of interstimulus intervals (ISIs) of 5 to 50 ms (*SI Appendix*, **Figure S1*A, B***). Thus, the differences in synaptic efficacy and reliability between S1 and PFC during the first action potential correlate with differential short-term plasticity, with the emphasize in S1 being on the first transmission process while in PFC facilitation emphasized the subsequent release.

Next, we investigated transmission at synapses between L2/3 and L5PNs, using minimal extracellular stimulation near somata of L2/3PNs (**Figure 1*A***). We found that also L2/3-L5PN synapses in PFC showed PPF (1.36, 1.14-1.88) as opposed to PPD in S1 (0.87, 0.72-0.99; **Figure 1*J, K***). We observed this over a range of ISIs up to 100 ms (*SI Appendix*, **Figure S1*C, D***). In addition, at PFC synapses the SD of the synaptic delays (0.46, 0.29-0.66) was again significantly larger than in S1 (0.25, 0.15-0.37; **Figure 1*L***). Furthermore, we found larger EPSC decay time constants (τ_decay_) in PFC compared to S1, possibly reflecting a contribution of different postsynaptic glutamate receptors (*SI Appendix*, **Figure S2*A, B***). Indeed, application of the N-methyl-D-aspartate receptor (NMDAR) antagonist 2-Amino-5-phosphonovaleriansäure (APV) removed the difference in τ_decay_ between PFC and S1 (*SI Appendix*, **Figure S2*C-F*)**. We extended the paired pulse experiments at L2/3-L5PN synapses in PFC to even older mice (P90-100) and to the mPFC (P21-26). We found that PPF persisted in the older age window (1.25, 1.14-1.41) and that also the synapses in mPFC showed PPF (1.14, 1.03-1.42; **Figure 1*K***).

Together these results from glutamatergic synapses onto L5PNs show significant differences in the synaptic delays, efficacy, reliability and short-term plasticity between PFC and S1. The results would be consistent with area-specific differences in the functional coupling distances between VGCCs and SVs at the presynaptic active zones.

### EGTA sensitivity of release is higher in PFC than in S1

To directly test for differences in the coupling distances, we investigated the sensitivity of release to synthetic Ca^2+^ chelators (**Figure 2**). The sensitivity of release to low millimolar concentrations of the kinetically slow Ca^2+^ chelator EGTA is a standard indicator of loose coupling (Eggermann et al., 2012). First, we performed recordings at pairs of L5PNs in mature PFC (**Figure 2*A, B***). After 10 min of stable baseline recordings, the presynaptic neuron was patched a second time and equilibrated for 30 min with 10 mM EGTA-containing pipette solution in the whole-cell configuration. In these paired recordings we found a significant effect of EGTA onto release in PFC (0.67, 0.63-0.73). In control recordings a temporary initial increase in EPSC amplitudes after whole-cell access to the presynaptic neuron was evident. Although L5PNs do not express large concentrations of mobile buffers (Helmchen et al., 1996; Tran and Stricker, 2018; Bornschein et al., 2019b), their wash-out might have contributed to the temporary run-up. On the other hand, run-up correlated with a temporary decline in R_s_ in some recordings (3 out of 6). Run-up effects normalized after 20 min at the latest (1.02, 0.75-1.20). In identical experiments in S1, we had previously found release to be insensitive to 10 mM EGTA (1.04, 0.80-1.10; control: 1.08, 0.86-1.31)(Bornschein et al., 2019b). For comparison data from the previous study were reanalyzed for the test period between 20 and 30 min. EPSC amplitudes in S1 were sensitive to the same concentration of the kinetically fast Ca^2+^ chelator BAPTA (test period 10-20 min: 0.22, 0.13-0.33). These data provide evidence that the synapses between L5PNs in mature PFC do indeed operate with loose coupling, in contrast to their counterparts in S1.

**Figure 2.**
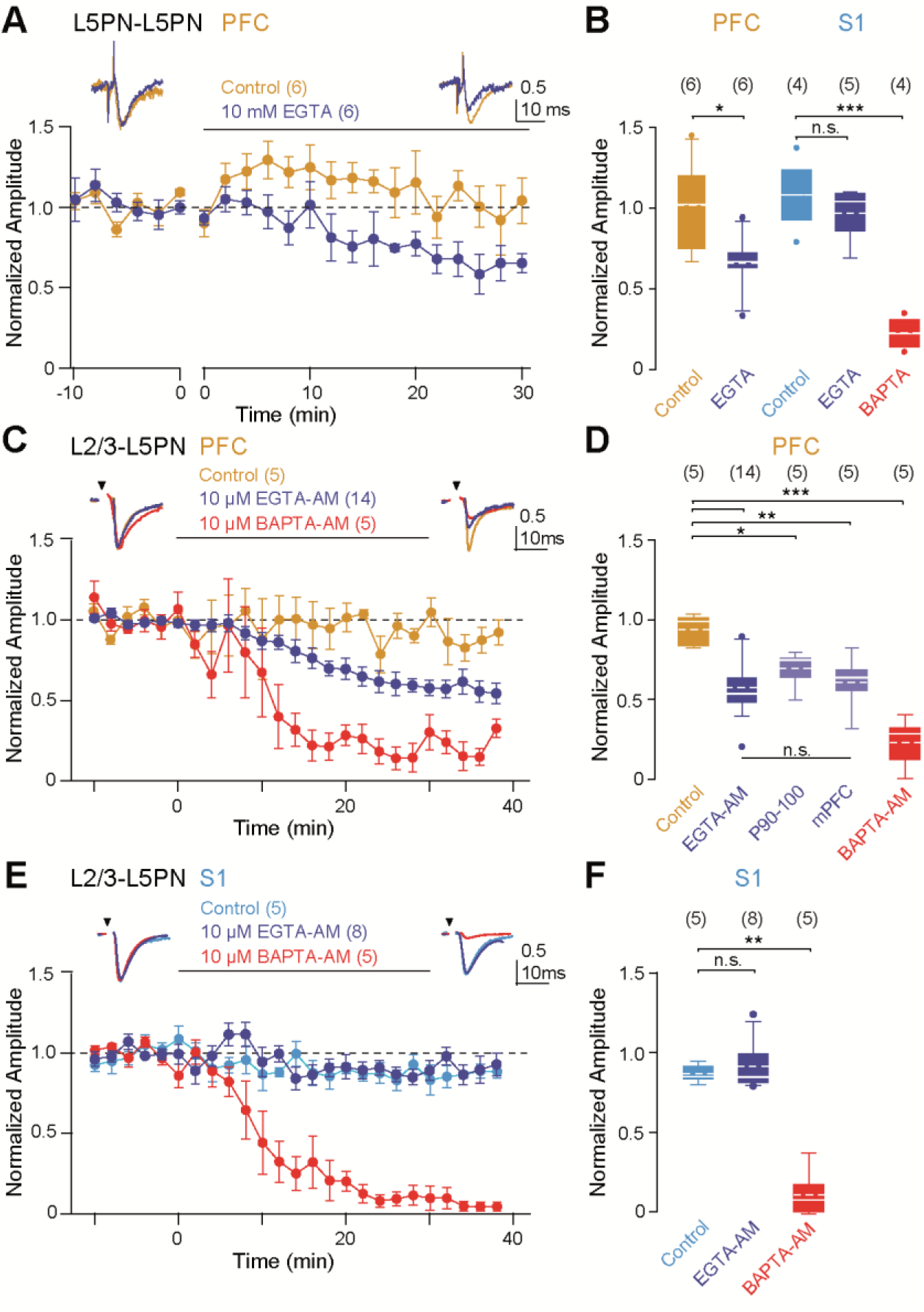
Area-specific differences in EGTA sensitivity. ***(A)***Averaged, baseline-normalized EPSC amplitudes (means ± SEMs, 2-min bins) recorded from L5PN-L5PN pairs in PFC with standard pipette solution (control, orange) and after re-patching the presynaptic neuron with a pipette solution containing 10 mM EGTA (blue). Insets: Average EPSCs from one pair during baseline (-10 to 0 min) and test period (20 to 30 min) for control and EGTA (averages normalized to baseline). ***(B)***Boxplot comparing the normalized EPSC amplitudes prior and following the application of EGTA and/or BAPTA with control recordings in L5PN pairs in PFC (*P=0.041, MWU) and S1 (***P<0.001, P=0.365, one-way ANOVA; Shapiro-Wilk normality test: P=0.597). S1 data are from ref. (Bornschein et al., 2019b). Note the significant EGTA effect. ***(C)***As in (***A***) but for L2/3-L5PN connections in PFC. Following baseline recordings (10 min), slices were perfused for 30 min (solid line) with ACSF containing either 10 μM EGTA-AM (blue), 10 μM BAPTA-AM (red), or only the solvent DMSO/Pluronic (control, orange), and thereafter rinsed for 10 min with ACSF (test period). Insets: Averaged EPSCs during baseline and test period. ***(D)***Summary of chelator-AM effects in PFC, mPFC and older PFC (P90-100; ***P<0.001, **P=0.009, *P=0.032, one-way ANOVA vs. control; Shapiro-Wilk normality test: P=0.666). Note the similar EGTA sensitivity of release in all PFC recordings (P=0.347, one-way ANOVA). ***(E)***As in (***C***) but for L2/3-L5PN connections in S1 (control in light blue). ***(F)***Summary of chelator-AM effects in S1 (**P=0.006, ANOVA on ranks). Note the absence of a significant EGTA effect in S1 (P=0.943, MWU).

We proceeded by comparing the EGTA sensitivity of release at L2/3-L5PN synapses between S1 and PFC, using extracellular stimulation and incubation with Ca^2+^ chelator-AM compounds (**Figure 2*C-F***) (Baur et al., 2015; Kusch et al., 2018). Following 10 min of stable baseline recordings, slices were perfused with ACSF that either contained the solvent DMSO/Pluronic alone (control) or 10 µM of dissolved EGTA-AM or BAPTA-AM. The fast BAPTA interferes with release irrespective of the coupling topography, thereby, providing a positive control for the proper functioning of the AM-method (Eggermann et al., 2012). EPSC amplitudes were quantified during a subsequent 10 min period with perfusion with normal ACSF. During this test period, BAPTA significantly reduced EPSC amplitudes both in PFC (0.28, 0.12-0.32; control: 0.99, 0.84-1.02) and in S1 (0.08, 0-0.18; control: 0.85, 0.83-0.92). On the other hand, EGTA significantly reduced EPSC amplitudes only in PFC (0.54, 0.47-0.67; 55% of control) but not in S1 (0.85, 0.81-1.04; 100% of control). For a better comparison of EGTA effects some recordings were performed in PFC and S1 derived from the same animal to rule out interindividual effects (see example recording in **Figure S3*A-C***).

In S1 we previously found that coupling switches from loose to tight during postnatal development between ∼P10 and ∼P20 (Bornschein et al., 2019b). To probe for a developmental retardation of this process in PFC, we extended the EGTA-AM experiments at L2/3-L5PN synapses to the older age-window of P90-100. Also, in these experiments we found a significant sensitivity of release to EGTA (0.76, 0.65-0.78; **Figure 2*D***; *SI Appendix*, **Figure S3*D***), indicating that loose coupling is a functionally persistent feature of synapses in PFC. Finally, we also extended the EGTA-AM experiments to synapses in mPFC and again found significant effects of EGTA on the EPSC amplitudes (0.65, 0.49-0.75; **Figure 2*D***; *SI Appendix*, **Figure S3*E***).

Together the results obtained with the slow Ca^2+^ chelator EGTA suggest that major glutamatergic synapses in mature PFC operate with loose coupling in contrast to tight coupling in the mature S1. The data further suggest that loose coupling is a developmentally persistent property of synapses in PFC. Finally, they indicate that the above functional differences between synapses in S1 and PFC originate from the differences in Ca^2+^ influx-release coupling.

### Release probabilities are similar in PFC and S1

To test whether the release probabilities (*p*_N_) of the differentially coupled release sites in PFC and S1 differ, we performed multiple probability fluctuation analysis (MPFA) (Clements and Silver, 2000; Brachtendorf et al., 2025) at different extracellular Ca^2+^ concentrations ([Ca^2+^]_e_; **Figure 3**) (Bornschein et al., 2019b). The parabolic MPFA-fits to variance-mean plots from recordings of pairs of L5PNs in PFC yielded a median *p*_N_ of 0.50 (0.37-0.55; **Figure 3*A, D***). This value was slightly but not significantly smaller than *p*_N_ of 0.66 (0.55-0.70), which we quantified previously for this connection in S1 (Bornschein et al., 2019b).

**Figure 3.**
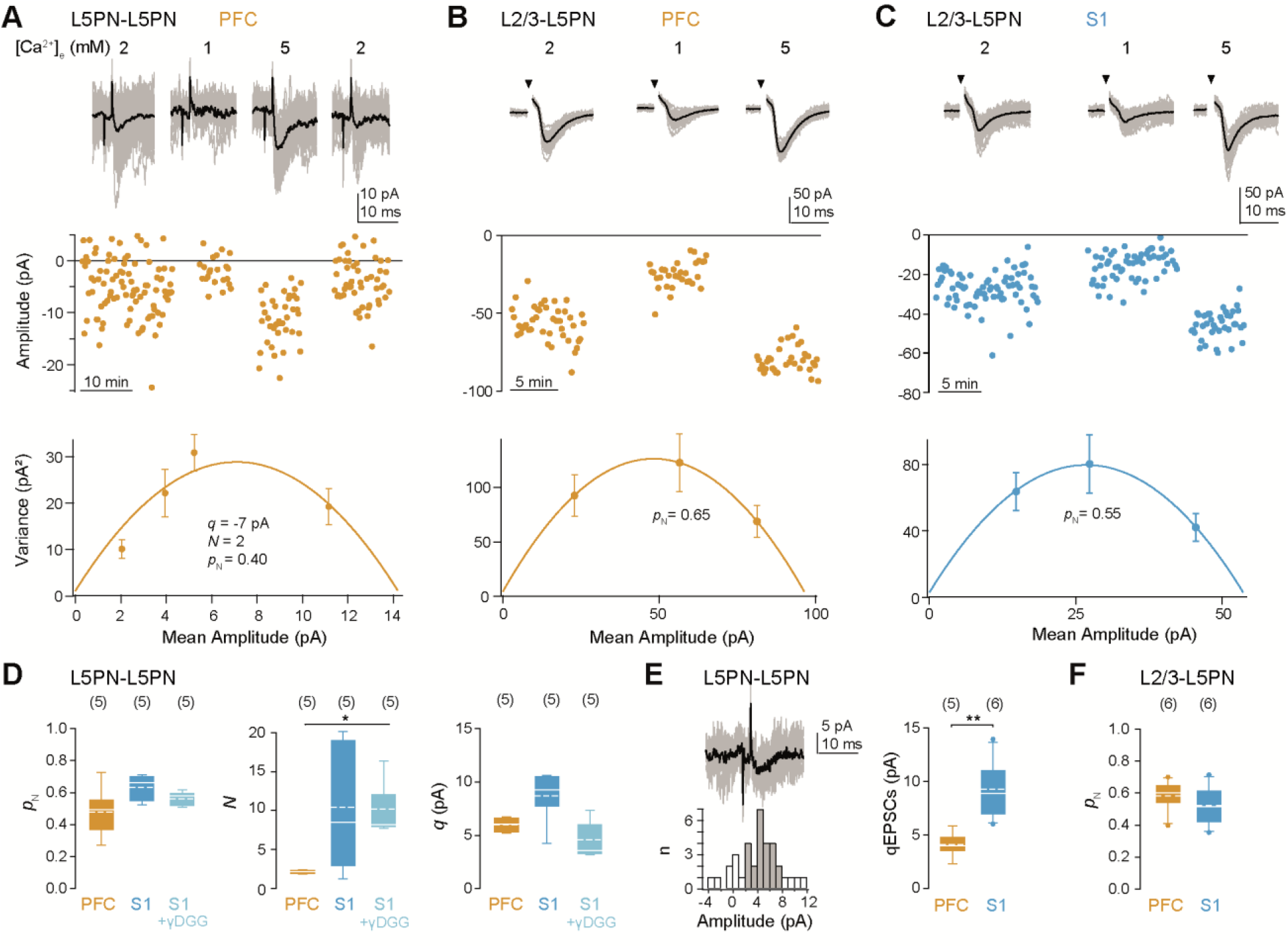
Quantal synaptic parameters in PFC and S1. ***(A)***MPFA of EPSC amplitudes recorded at the indicated [Ca^2+^]_e_ from a pair of L5PNs in PFC. Top: EPSCs (gray, average in black) recorded in 1, 2 and 5 mM [Ca^2+^]_e_. Middle: Plot of EPSC amplitudes over time. Bottom: Corresponding mean-variance plot fitted with a parabola estimating the quantal parameters of release. *p*_N_ is for 2 mM [Ca^2+^]_e_. ***(B)***As in (***A***), but for a L2/3-L5PN connection in PFC. Recordings were made in the presence of 0.25 mM Kyn and 50 µM APV. ***(C)***As in (***B***), but in S1. ***(D)***Summary of quantal release parameters in L5PN pairs from PFC and S1 and in S1 with 1-2 mM γDGG (S1 data from (Bornschein et al., 2019b); P=0.121, *P=0.048, P=0.054, ANOVA on ranks). ***(E)***Left: Example of evoked qEPSCs (top) and the corresponding amplitude histogram (bottom, gray range was used for calculating the qEPSC amplitude). Right: Summary of qEPSC amplitudes in PFC and S1 (**P=0.004, MWU). ***(F)***Summary of *p*_N_ values in L2/3-L5PN connections from PFC and S1 (P=0.485, MWU).

We also quantified *p*_N_ at L2/3-L5PN synapses, both in PFC and in S1, using MPFA with extracellular stimulation (Baur et al., 2015). We found that *p*_N_ was as high as at the L5PN-L5PN synapses in both areas with values (PFC: 0.60, 0.51-0.66; S1: 0.52, 0.40-0.64) being not significantly different between areas (**Figure 3*B, C, F***). Interestingly, in our previous study (Bornschein et al., 2019b) we found that the developmental switch from loose to tight coupling at L5PN synapses in S1 also had no significant impact on *p*_N_.

From MPFA in paired recordings it is possible to estimate the number of release sites (*N*) and the quantal size (*q*), additionally. *N* was significantly smaller in PFC than in S1 being only 2.1 (1.9-2.3) rather than 8 (3-19). *q* tended to be smaller in PFC (6 pA, 5-7 pA) than in S1 (9 pA; 8-11 pA) and the difference became significant when evoked quantal EPSCs recorded in low [Ca^2+^]_e_ were compared (PFC: 4 pA, 4-5 pA; S1: 9 pA, 7-11 pA). We consider it unlikely that postsynaptic receptor saturation or desensitization influenced the determination of quantal parameters since experiments with the competitive low-affinity glutamate receptor antagonist γ-DGG (γ-D-glutamylglycine) (Chanda and Xu-Friedman, 2010) performed at the S1 synapses had revealed no signs of this (*p*_N_=0.58, 0.52-0.60; *N*=8, 8-12; **Figure 3*D***). Due to the smaller size of the EPSCs, testing on the PFC synapses was not possible. Estimating EPSC amplitudes (EPSC = *N p*_N_ *q*; PFC, 6 pA; S1, 48 pA) and failure rates (F = (1-*p*_N_)^*N*; PFC, 0.25, S1, 0.0001) from the quantal parameters yielded values similar to those recorded (**Figure 1**). The determined parameters were therefore well suited to explain the area-specific differences in synaptic efficacy during individual action potentials.

### Presynaptic Ca^2+^ signals are similar in PFC and S1

For a thorough interpretation of the above results knowledge about the presynaptic Ca^2+^ dynamics in the different synapses is required. We performed dual-dye two-photon Ca^2+^ imaging (Sabatini et al., 2002) to quantify volume-averaged Ca^2+^ signals at presumed presynaptic boutons located on axon collaterals of L5PNs in PFC and in S1. The imaged presynaptic boutons most likely connect to neighboring pyramidal cells, as connectivity between L5PNs in layer 5A is high (Feldmeyer, 2012). Nevertheless, a small proportion of other postsynaptic targets, such as interneurons, cannot be ruled out. We quantified single action potential-mediated elevations in green over red fluorescence signals (ΔG/R) that were converted to increases in intracellular calcium (Δ[Ca^2+^]_i_) (**Figure 4**) (Bornschein et al., 2019b). We found that neither the Ca^2+^ transients nor the decay time constants were significantly different between the cortical areas (**Figure 4*A-E***; PFC: G/R=0.25, 0.23-0.31; Δ[Ca^2+^]_i_=279 nM, 244-453 nM; τ_decay,1_=9 ms, 6-19 ms; τ_deay,2_=91 ms, 75-120 ms; S1: G/R=0.29, 0.25-0.39; Δ[Ca^2+^]_i_=238 nM, 208-303 nM; τ_decay,1_=9 ms, 3-15 ms; τ_decay,2_=86 ms, 63-212 ms). The quantification of absolute basal [Ca^2+^]_i_ was limited by the *K*_D_ of Fluo5F (439 nM), which slightly underestimated basal [Ca^2+^]_i_ values in comparison to quantification with OGB1 (K_D_=166 nM, **Figure S4**). Nevertheless, relative comparison of basal [Ca^2+^]_i_ yielded no significant differences between PFC (30 nM, 21-34 nM) and S1 (22 nM, 13-38 nM; **Figure 4*F***).

**Figure 4.**
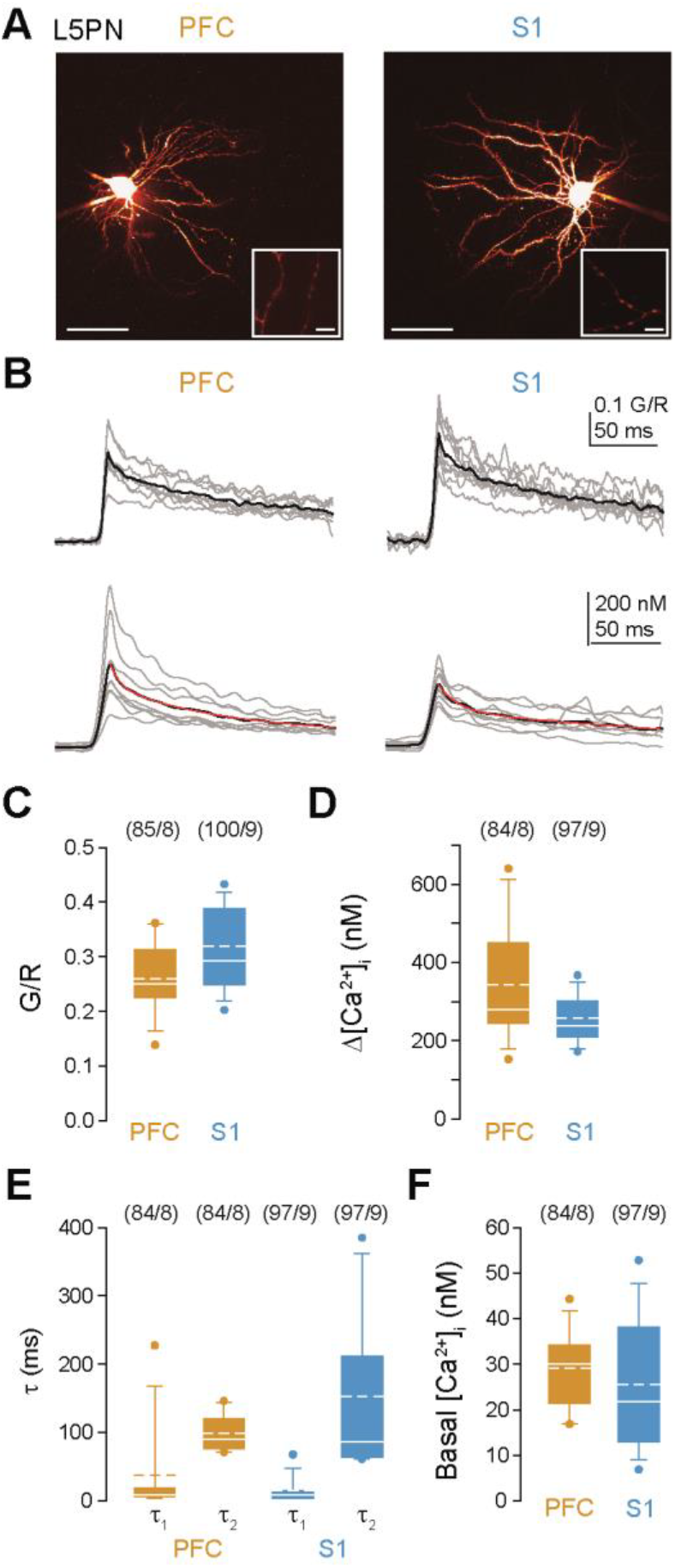
Presynaptic Ca^2+^ transients are similar in PFC and S1. ***(A)***Two-photon images of L5PNs in PFC (left) and S1 (right) filled with Fluo-5F and Alexa 594 (scale bar 50 µm). Insets: Magnifications showing presynaptic boutons from which Ca^2+^ transients are recorded (scale bar 5 µm). ***(B)***G/R signals (top) evoked by single APs in individual cells (gray, average of 6-15 boutons each, grand average in black) and their conversion to Ca^2+^ increases (Δ[Ca^2+^]_i_, bottom). **(*C-F*)** Summary of G/R signals (***C***; P=0.112, MWU; numbers of boutons and cells in brackets), Δ[Ca^2+^]_i_ (***D***; P=0.361), decay time constants τ_1_ and τ_2_ (***E***; P=0.532, 0.962), and basal Δ[Ca^2+^]_i_ (***F***; P=0.597) in PFC and S1. Note, that there is no significant difference in presynaptic Ca^2+^ transients between PFC and S1.

The similarity in the presynaptic Ca^2+^ transients of boutons in PFC and S1 suggests that differences in the presynaptic Ca^2+^ influx are unlikely to be the cause for the differential Ca^2+^ chelator effects. Rather, they support the view that these differences result from loose coupling in PFC and from tight coupling in S1.

### Estimate of the coupling distance in PFC verify loose microdomain coupling in PFC

To derive a quantitative picture of the coupling topography in L5PN synapses in PFC we used numerical computer simulations that were based on and constrained by the experimental data. First, we simulated the measured Ca^2+^ transient with the indicator dye present in the simulation (**Figure 5*A***). Subsequently, the indicator dye was removed and a release sensor model for Synaptotagmin-1-triggered release from L5PN boutons was included in the simulations (**Figure 5*B, C***; *SI Appendix*, **Figure S5**) (Bornschein et al., 2025). VGCCs were assumed to form a ring like structure around a vesicle (**Figure 5*D***), similar to the release site topography in young S1 (Bornschein et al., 2019b). Release rates as reported by the sensor model were integrated over time to yield the *p*_N_ values. The coupling distance between the sensor and the ring of VGCCs as well as the number of VGCCs forming the ring were iteratively varied until the simulation correctly predicted the experimental *p*_N_ values from MPFA obtained under control conditions and in the presence of EGTA. By this procedure we quantified an average coupling distance of 48-51 nm between a microdomain of several VGCCs and the release sensor (**Figure 5*B***; *SI Appendix*, **Figure S5**), similar to the estimate for these synapses in immature S1 (Bornschein et al., 2019b). Thus, a microdomain model similar to immature S1 is suitable to predict the data from mature PFC, whereas the same synapses in mature S1 operated with Ca^2+^ signaling nanodomains in which only 1-3 VGCCs at distances of 11-16 nm trigger fusion (Bornschein et al., 2019b; Bornschein et al., 2025).

**Figure 5.**
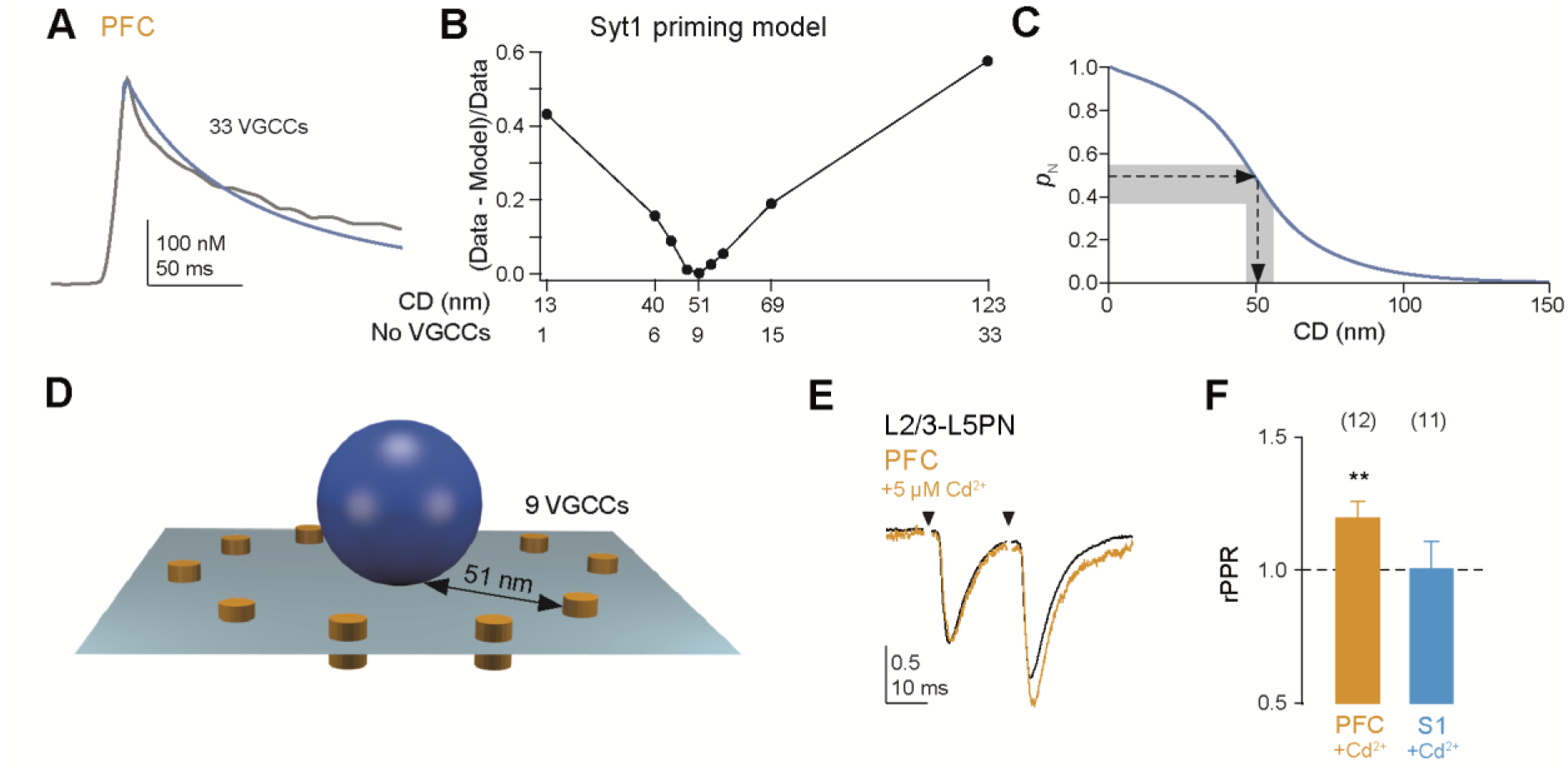
Model predicts microdomain coupling at PFC synapses. ***(A)***Fit of the model (blue line) to the measured average Ca^2+^ signal (***A***; gray line) of L5PN boutons in PFC. The fit was obtained with the indicated number of open VGCCs. ***(B)****p*_N_ values were simulated with the Syt1 priming model (Bornschein et al., 2019b; Bornschein et al., 2025) for control conditions and in the presence of EGTA with different VGCC numbers yielded different coupling distances (CD) in the model. The simulated *p*_N_ were subtracted from and normalized to the experimental values, i.e. a value close to zero indicates best agreement between simulation and experiment. This was obtained with 9 open VGCCs at a CD of 51 nm (see **C**). ***(C)***Simulated *p*_N_ for 9 open VGCCs under control conditions at increasing CDs (blue line). The experimental *p*_N_ value (median ± IQR, dashed line, gray range, **Figure 3D**) was reproduced at a CD of 51 nm (48-55 nm). ***(D)***Illustration of the ring model used in **B** and **C** with a CD of 51 nm between 9 VGCCs (orange) and the release sensor of a vesicle (blue). ***(E)***Examples of averaged normalized paired pulse recordings at 50 Hz at L2/3-L5PN synapses before (black) and after (orange) application of 5 µM Cd^2+^ in PFC. ***(F)***Bar graph quantifying normalized PPRs after Cd^2+^ application in PFC and S1 (**P=0.005, P=0.667, paired t-Test; Shapiro-Wilk normality test: P=0.514). Note that Cd^2+^ increased the PPR in PFC.

To experimentally test whether overlapping Ca^2+^ domains from several VGCCs trigger release in PFC (microdomain), we analyzed the effects of the unspecific VGCC blocker Cd^2+^ onto the PPR. If fusion is triggered by such VGCC microdomains, application of a subsaturating concentration of Cd^2+^ will increase the PPR, while it will leave PPR unaffected if only a single or few VGCCs trigger release (Hefft et al., 2002; Bucurenciu et al., 2008; Scimemi and Diamond, 2012; Baur et al., 2015; Bornschein et al., 2019b). In paired pulse experiments, at L2/3-L5PN synapses in PFC we indeed found a significant increase in the PPR following the application of 5 µM Cd^2+^ (rPPR=1.20±0.06), whereas the average PPR remained unaffected in S1 (rPPR=1.01±0.10; **Figure 5*E, F***). Thus, our data and simulations suggest that the nanotopography of PN synapses in mature PFC is reminiscent of the same synapses in young S1.

## Discussion

Our results provide evidence that the same archetypes of glutamatergic synapses show area-specific differences in their functional release site nanoarchitectures in the mature neocortex. They suggest loose microdomain coupling as a synaptic correlate of higher order neocortical functions in PNs.

We did neither morphologically nor based on spiking patterns differentiate further between PN subtypes within a given layer. However, within a cortical area (S1 or PFC) we did not find a difference between glutamatergic synapses from L2/3 onto L5PNs and L5PN to L5PN synapses, neither with regard to the EGTA sensitivity of release nor with regard to *p*_N_. In particular, we found homogeneous results and similar variability in both, the examined connections in the PFC and in S1, with no discernible clustering in the data that would indicate stimulation of different cell populations. These findings suggest that excitatory inputs to L5PNs exhibit similar properties (PPR, *p*_N_, CD) irrespective of whether they originate in L2/3 or in neighboring PNs in L5A. However, we do see significant differences between synapses in the different cortical areas S1 and PFC. Thus, intra-area specific differences in morphology and spiking patters among PNs appear to be not reflected on the level of their synapses.

The differences in coupling gave rise to larger variability of synaptic delays (SD_Delay_) in PFC compared to S1 and were associated with differences in synaptic reliability and efficacy, whereas the *p*_N_ values were not significantly different between areas. A major determinant of *p*_N_ is the size of the Ca^2+^ signal at the release sensor. Accordingly, tightening of coupling can increase *p*_N_ (Eggermann et al., 2012; Baur et al., 2015; Bornschein and Schmidt, 2019). However, large Ca^2+^ elevations at the release sensor can also be obtained with Ca^2+^ microdomains. At L5PN-L5PN synapses in S1 the developmental switch from microdomain to nanodomain coupling had no effect on *p*_N_ (Bornschein et al., 2019b). At the calyx of Held (Iwasaki and Takahashi, 2001; Taschenberger et al., 2002; Fedchyshyn and Wang, 2005; Koike-Tani et al., 2008; Nakamura et al., 2015) and at cerebellar basket cell to Purkinje cell synapses (Chen et al., 2024) *p*_N_ even decreased during development, whereas coupling got tightened. The decrease in *p*_N_ was compensated by an increase in *N* at these synapses, which increased their reliability and efficacy during development. We found *N* to be larger in S1 than in PFC. Thus, it appears that differences in coupling are typically associated with other characteristic changes in the release parameters. Overall, this suggests that reliable CNS synapses switch from an initial state with high plasticity to a matured state with high reliability. On the other hand, PN synapses in the PFC appear to remain in a more juvenile state that favors plasticity over reliability.

Nanodomain coupling was also found in the peripheral nervous system, in particular at retinal (Singer and Diamond, 2003; Jarsky et al., 2010) and auditory (Moser and Beutner, 2000; Brandt et al., 2005) ribbon type synapses and at the neuromuscular junction (Harlow et al., 2001; Shahrezaei et al., 2006). These synapses have highly specialized properties and appear to be optimized for very reliable transmission and, in the case of ribbon synapses, also for high-frequency coding of sensory information (reviewed in Matthews and Fuchs, 2010; Eggermann et al., 2012). Thus, it appears that synapses in the sensory pathways, in particular those engaged in reliable high-frequency coding of sensory information, both in the periphery and in the lower processing stages of the CNS, up to primary sensory cortices, operate with nanodomain coupling. In the executing motor pathway, the neuromuscular junction uses nanodomain coupling and, as recent results from our group suggest, also PNs in the primary motor cortex (Yarim et al., *in preparation*). It is tempting to speculate that complete loops from or to the primary cortices to their peripheral target organs operate with nanodomains. Microdomain coupling, on the other hand, appears to come into play only if integration of information from multiple sources and plasticity are the main focus, as at certain synapses in PFC (this study) or hippocampus (Vyleta and Jonas, 2014).

The microdomain was assumed to be formed by a ring-like structure of VGCCs around a vesicle (**Figure 5*D***). This topography was chosen because such a microdomain was found to best predict the experimental data of transmitter release from PNs in young S1 (Bornschein et al., 2019b). Other previously described distributions of VGCCs suitable to reproduce release data cover random distributions of VGCCs (Scimemi and Diamond, 2012), VGCC clusters (Meinrenken et al., 2002; Nakamura et al., 2015; Rebola et al., 2019), and exclusion zones (Keller et al., 2015; Rebola et al., 2019). In the early S1, all of these models predicted a higher EGTA sensitivity of the microdomain, however, these models provided a poorer fit to the full set of the experimental data than the ring-like structure (Bornschein et al., 2019b). Since the experimental data from PNs in the mature PFC were similar to those in young S1, these other microdomain models were not tested explicitly here.

Differences in the sensitivity of release to low to moderate concentrations of EGTA (≤ 30 mM) are a standard indicator of differences in the coupling distance (e.g. Adler et al., 1991; Bucurenciu et al., 2008; reviewed in Eggermann et al., 2012; Vyleta and Jonas, 2014; Kusch et al., 2018; Bornschein et al., 2019b). *p*_N_ is determined by the size of the Ca^2+^ signal at the release sensor and the binding kinetics and affinity of the sensor. The former in turn is determined by the details of the Ca^2+^ influx and the diffusional coupling distance between the VGCCs and the sensor. The similarity of Ca^2+^ signals between synapses in PFC and S1 (**Figure 4**) indicates that Ca^2+^ influx is similar between boutons, although more subtle differences in the influx kinetics may have remained undetected in these volume-averaged signals. Regarding sensor affinity, results in a previous study indicate that differences in EGTA sensitivity show differences in coupling rather than sensor affinity even if *k*_on_ of the sensor and its affinity should differ as much as ten-fold, which appears to be an unlikely scenario given that even the two major isoforms of Synaptotagmin that trigger synchronous release differ by less than a factor of three to four in their affinity (Bollmann et al., 2000; Schneggenburger and Neher, 2000; Bornschein et al., 2025). Finally, the increase in the PPR induced by the application of Cd^2+^ further supports our conclusion of microdomain coupling in the PFC synapses (Scimemi and Diamond, 2012). Thus, although the absolute EGTA sensitivity is influenced by different factors, which necessitates data-constrained models for quantitative comparisons, the general sensitivity of release to EGTA indicates loose coupling.

The differences we observed in short-term plasticity suggest that S1 emphasized the first transmission process, whereas in PFC facilitation emphasized the subsequent release. The mechanisms of short-term plasticity are complex and include the regulation of *p*_N_, the speed of vesicle replenishment and the recruitment of release sites (Neher and Brose, 2018; Schmidt, 2019; Neher, 2023; Brachtendorf et al., 2025). Classically, *p*_N_ was considered as the major determinant of short-term plasticity (e.g. reviewed in Zucker and Regehr, 2002; Feldmeyer and Radnikow, 2009). More recently other factors, including the number of occupied release sites, their replenishment or an increase in their occupancy, or the expression of endogenous Ca^2+^ buffers have been considered as more important determinants of short-term plasticity (Rozov et al., 2001; Blatow et al., 2003; Felmy et al., 2003; Matveev et al., 2004; Bornschein et al., 2013; Miki et al., 2016; Doussau et al., 2017; Jackman and Regehr, 2017; Neher and Brose, 2018). Irrespective of the detailed mechanism, we consider the differential emphasize on the available presynaptic efficacy between PFC and S1 as a signature of higher plasticity and flexibility in the processing of information in PFC compared to S1.

Short-term plasticity changes during postnatal development at different cortical PN connections without alterations in *p*_N_ (Reyes and Sakmann, 1999; Bornschein et al., 2019a). For L5PN to L5PN connections these differences were found to result from the maturation of an intermediate replenishment vesicle pool (Bornschein et al., 2019a). Such pool maturation may also underlie the elimination of layer-specific differences in short-term plasticity between L2/3-L5B and L5B-L5B synaptic connections that were evident in young rats but eliminated during the first weeks of postnatal development (Reyes and Sakmann, 1999). Since postnatal development follows the same time-course in mouse PFC and S1 and occurs predominantly within the first two weeks after birth, significant layer-specific differences in PN synapses are unlikely in both areas in our experimental time window (Kroon et al., 2019). Consistently, we found similar PPRs at L2/3-L5PN synapses and L5PN-L5PN synapses in both areas, with facilitation in PFC and depression in S1, irrespective of the presynaptic PN synapse type.

Two other previous studies on L5APN (Frick et al., 2008) and L5BPN (Markram et al., 1997) connections concluded that these synapses operate with high release probability, which nicely agrees with our previous (Bornschein et al., 2019b) and current results. It is remarkable that we did not even detect any differences between the L2/3-L5PN and L5PN-L5PN synapses within a given cortical area. Overall these results from different studies (Markram et al., 1997; Reyes and Sakmann, 1999; Frick et al., 2008; Bornschein et al., 2019a; Bornschein et al., 2019b) may indicate that variability on the synaptic level between PNs of a given area is not pronounced. In order to further substantiate this, we determined the relative variability in median EPSC amplitudes to test whether there is a higher variability of recorded cell types in PFC compared to S1. The relative MAD (median absolute deviation) of EPSC amplitudes was 0.46 in PFC and 0.50 in S1. The similarity in these values argues against higher cell-type variability in PFC compared to S1.

Our results have implications for long-term plasticity at neocortical PN synapses. Both, long-term potentiation (LTP) and depression (LTD) have a presynaptic locus at these synapses and likely involve changes in *p*_N_, but the final steps that alter release are not well understood (Feldman, 2009; Castillo, 2012; Feldman, 2012). A recent study on L5PN synapses in S1 showed an increase in the number of primed vesicles as a major mechanism of LTP (Weichard et al., 2023). PN synapses in S1 are already tightly coupled and therefore a further decrease in the coupling distance is an unlikely mechanism of LTP. However, this does not exclude that in S1 LTD induction breaks tight coupling and vice versa that LTP in PFC involves tightening of coupling. In addition, in PFC we found an increased contribution of NMDAR on the postsynaptic site. Thus, these synapses have increased potential for plasticity on the pre- and the postsynaptic site. During induction of long-term plasticity, the postsynaptic site signals back to the presynaptic site via retrograde messengers (Regehr et al., 2009) and loose coupling will provide the substrate for enhanced regulatory capacity (Vyleta and Jonas, 2014).

In general, loose coupling provides more flexibility to regulate [Ca^2+^]_i_ at the release sensor (Vyleta and Jonas, 2014). Prefrontal networks carry out higher-order computations that transform sensory information and memory to perform flexible output such as delayed response, inference and planning (Narayanan et al., 2025). Our findings suggest that loose coupling provides a synaptic foundation for such flexible neocortical network functions.

## Methods

### Slice preparation

C57BL/6J mice at P21-26 of either sex were used in this study. All experiments were performed in accordance with the guidelines for the welfare of experimental animals issued by the European Communities Council Directive (2010/63/EU) and with the German Protection of Animals Act (Tierschutzgesetz). Mice were bred in the animal facility of the Medical Faculty of Leipzig University and were housed in individually ventilated cages in a specific pathogen free environment and in a 12h/12h light dark cycle with access to food and water ad libitum. Experiments were approved by the animal welfare office of the University Medical Center, Leipzig, as well as the local governmental authorities (Landesdirektion Leipzig, registration numbers T10/20, T05/21-MEZ).

C57BL/6J mice at P21-26 and P90-100 of either sex were decapitated under deep Isoflurane (Curamed) inhalation anesthesia. The brain was excised rapidly and placed in cooled (0-4°C) artificial cerebrospinal fluid (ACSF) containing (in mM): 125 NaCl, 2.5 KCl, 1.25 NaH_2_PO_4_, 26 NaHCO_3_, 1 MgCl_2_, 2 CaCl_2_, and 20 glucose, equilibrated with 95% O_2_ and 5% CO_2_ (pH 7.3-7.4). Coronar neocortical slices (150-250 μm thick) were cut from the lateral PFC, medial PFC (mPFC) or S1 region (**Figure 1*A***) with a vibratome (HM 650 V, Microm). For some paired recordings between L5PNs and for calcium imaging experiments parasagittal slices (200 µm) were prepared from PFC region. Slices were incubated for 30 min at 35°C and subsequently stored at room temperature. For experiments, slices were transferred to a recording chamber and continuously perfused with ACSF (2-3 ml per min, supplemented with 10 μM (-)-bicuculline methiodide (Tocris) at 31-33°C. Unless stated otherwise, chemicals were from Sigma-Aldrich.

### Electrophysiological recordings

Patch-clamp recordings from L5PNs located in the upper layer 5 (L5A in S1) were established according to the criteria described in detail in our previous work on this connection in S1 (Bornschein et al., 2019b; Bornschein et al., 2025). Presynaptic neurons were stimulated extracellularly in upper layer 2/3 (L2/3-L5PN connections) straight above the patched L5PN or in on-cell mode in L5A right next to the postsynaptic cell (L5PN-L5PN connections; **Figure 1**).

Patch pipettes were prepared from borosilicate glass (Hilgenberg) with a PC-10 puller (Narishige) and had final resistances of 6-8 MΩ when filled with the following standard pipette solution (in mM): 150 K-gluconate, 4 NaCl, 3 MgCl_2_, 3 Na_2_ATP, 0.3 NaGTP, 0.05 EGTA, 10 KHEPES, dissolved in purified water. The pH was adjusted to 7.3 with KOH. Recordings were performed under optical control (BX51WI, Olympus), using an EPC10/2 amplifier and Patchmaster software (version v2x90.2, HEKA). EPSCs were recorded in the whole-cell configuration at a holding potential (V_hold_) of -90 mV (online corrected for a liquid junction potential of 16 mV), filtered at 5 kHz and sampled at 10 kHz. Series resistance (R_s_) was continuously compensated to a fixed value between 10 and 15 MΩ. The average uncompensated R_s_ was 18±1 MΩ in L5PN-L5PN paired recordings (n=41 cells, mean ± SEM) and 23±1 MΩ in L2/3-L5PN recordings (n=133 cells). Holding current (I_hold_) was also monitored continuously and was <180 pA (-134±16 pA in L5PN-L5PNs; -168±11 in L2/3-L5PNs). Typically, recordings were excluded if R_s_ exceeded 30 MΩ and I_hold_ fell below -300 pA. In paired recordings, presynaptic L5PNs were stimulated in on-cell configuration (200-500 mV, 1-2 ms).

In the chelator wash-in experiments, presynaptic neurons were repatched with a pipette solution supplemented with 10 mM EGTA (K-gluconate concentration was reduced to 135 mM to adjust osmolarity) and whole-cell configuration was established to allow EGTA perfusion of the presynaptic neuron. EPSCs were evoked every 10 s for at least 30 min (**Figure 2, S3**). Averaged EPSC amplitudes were calculated before (≥10 min baseline) and 20-30 min after repatching (test period). For recordings from L2/3-L5PN connections ACSF-filled patch pipettes were placed close to the somata of PNs in L2/3 and extracellular stimulation was performed using an ISO-Stim 01 DPI (NPI electronics). Minimal stimulation (0.5 to 5 V) was used and yielded EPSC amplitudes of ∼40 pA (47±3 pA, n=131 pairs), making it likely that we investigated transmission mainly between single or few L2/3PNs and L5PNs in these experiments. Following 10 min of stable recordings in standard ACSF (baseline), the bath solution was exchanged for ACSF containing either 0.1% DMSO and 0.01% Pluronic (control solution), 10 µM EGTA-AM or 10 µM BAPTA-AM in DMSO/Pluronic for 30 min (incubation period) and thereafter replaced by standard ACSF again for another 10 min (test period). Averaged EPSC amplitudes were calculated during baseline and test period. The time courses of chelator effects on EPSC amplitudes were analyzed by binning EPSC amplitudes within 2 min intervals of recording time to average amplitudes. Chelator effects were expressed as averaged EPSC amplitudes during test period normalized to the corresponding baseline values.

VGCCs were inhibited by bath-application of ACSF containing a subsaturating concentration of 5 µM CdCl_2_ (**Figure 5*E, F***). NMDA receptors were blocked by the selective antagonist 2-Amino-5-phosphonovaleriansäure (APV, 50 μM; *SI Appendix*, **Figure S2**). For assessing the effects of the individual blockers, EPSCs were recorded in ACSF for at least 10 min (10 s intervals), subsequently the blocker was perfused and the blocker effects were quantified after a stable block had been established (at least 5 min test period).

### Quantification of quantal synaptic parameters

Quantal synaptic parameters were estimated from parabolic fits to mean-variance (MV) plots of EPSC amplitudes recorded at different [Ca^2+^]_e_ (1, 2, and 5 mM, ≥30 repetitions per concentration; [Mg^2+^]_e_ was correspondingly adjusted to 2, 1, 0 mM, respectively), assuming binominal release statistics (Clements and Silver, 2000; Scheuss et al., 2002; Silver, 2003)(**Figure 3**). The recordings always started in 2 mM [Ca^2+^]_e_. The variance of EPSCs (σ^2^) was calculated according to ref. (Scheuss et al., 2002), plotted against the averaged amplitudes (I), and fitted by a parabola of the form:

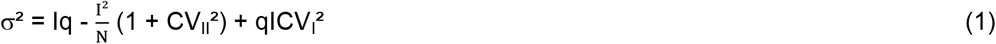

where *q* is the quantal size, *N* a binominal parameter, and CV_I_ and CV_II_ the coefficients of intrasite and intersite quantal variability, assumed to be 0.3 (Clements and Silver, 2000). For CV assumptions, that deviate from the standard value of 0.3, deviations in the calculated *p*_N_ values of less than 10% are to be expected (Schmidt et al., 2013; Bornschein et al., 2019b). The fits were constrained to pass through zero. The variance of the variance was calculated according to ref. (Meyer et al., 2001). Stationarity of EPSC amplitudes was established when, after full exchange of the bathing solutions (≥5 min), EPSC amplitudes no longer showed a tendency to increase or decline. For MV analysis in L2/3-L5PN connections, the bath solution was supplemented with the competitive AMPA receptor antagonist kynurenic acid (Kyn, 0.25 mM), which relieves their desensitization and saturation (Neher and Sakaba, 2001) and with 50 µM APV to prevent NMDAR activation. Due to the small size of EPSCs in L5PN-L5PN connections no blocker was used in these recordings. The size of qEPSCs was quantified from evoked miniature EPSCs recorded in low [Ca^2+^]_e_ (1 mM) and high [Mg^2+^]_e_ (2 mM).

### Ca^2+^ imaging

Action potential-evoked (current injection of 1-4 nA for 1-2 ms) fluorescence changes were recorded at boutons located on axon collaterals of L5PNs as described previously (Bornschein et al., 2019b; Bornschein et al., 2025). L5PNs were filled with EGTA-free, Fluo-5F (200 µM, Invitrogen) and Alexa-594 (50 µM, Molecular Probes) containing pipette solution via somatic whole-cell patch-pipettes (**Figure 4**). Basal [Ca^2+^]_i_ values in S1 were additionally determined by OGB-1 (100 µM, Invitrogen; **Figure S4**). Cells were dialyzed for at least 20 min to yield sufficient equilibration with the dyes. Volume averaged fluorescence signals were recorded in point mode at 500 kHz temporal resolution and subsequently binned to 500 Hz, using a custom-build two-photon microscope based on a Fluoview-300 scanner (Olympus), a 60x/0.9 N.A. objective, a mode-locked Ti:sapphire laser (Tsunami, Newport-Spectra Physics, set to a center wavelength of 810 nm), and a Pockels cell (350-80 KD*P, Conoptics). The fluorescence background was determined in a subsequent point mode recording performed near the boutons under investigation. On average 11 (but at least 3) boutons per L5PN were recorded and averages per cell were calculated to include a large number of putative presynapses. The fluorescence signals were filtered (HC647/75, Semrock HC525/50, 720-SP, AHF), detected by two external PMT modules (H7422-40, Hamamatsu; PMT-02M/PMM-03, NPI electronics) monitoring red and green epifluorescence, respectively at fixed PMT voltages, and digitized with the Fluoview system. The Ca^2+^-dependent green fluorescence was normalized to the Ca^2+^-insensitive red fluorescence and expressed as background-corrected ΔG/R (Sabatini et al., 2002). ΔG/R signals were converted to changes in [Ca^2+^]_I_ based on an *in vitro* quantification of the *K*_D_ of Fluo-5F (439 nM) and OGB-1 (166 nM), respectively, in our pipette solution, adjusted with CaEGTA and K_2_EGTA (100 mM stock solutions, 10 mM HEPES added) to contain different free Ca^2+^ concentrations that were calculated with the MaxChelator (https://somapp.ucdmc.ucdavis.edu/pharmacology/bers/maxchelator/CaMgATPEGTA-NIST.htm). R_min_ and R_max_ values were measured after each successful recording in sealed patch pipettes that were placed in the slices near the recorded cells. The pipettes contained the EGTA-free, Fluo-5F- and Alexa-594-containing pipette solution with either 0 mM CaCl_2_ and 10 mM K_2_EGTA (zero Ca^2+^ solution) or 20 mM CaCl_2_ (high Ca^2+^ solution).

### Modeling

The kinetic gating scheme of the VGCCs and the reaction schemes for Ca^2+^ binding to buffers and transmitter release were converted to ordinary differential equations (ODEs). The system of ODEs was numerically solved using “NDSolve” or “NDSolveValue” of Mathematica 14 (Wolfram) as described previously (Schmidt et al., 2013; Bornschein et al., 2019b; Bornschein et al., 2025). Spatial resolution was achieved by placing the ODEs in concentric hemi-shells (1 or 2 nm thickness) that were coupled via diffusion.

The presynaptic action potential was modeled with a half-duration of 0.2 ms, τ_rise_ of 0.1 and τ_decay_ of 0.2 ms (Borst et al., 1995). Voltage dependent activation of VGCCs was simulated using the gating scheme for P/Q-type channels, a single channel conductance of 2.5 pS, and a Ca^2+^ equilibrium potential of 130 mV (Li et al., 2007). The model included ATP as mobile buffer and immobile endogenous buffers with a buffer capacity (κ_E_) of 25 (Tran and Stricker, 2018). Model parameters are given in **Table S1** (*SI Appendix*).

The model for each channel consisted of five closed (C) and one open (O) state. The transition between the first five steps was voltage dependent and the last step voltage independent:

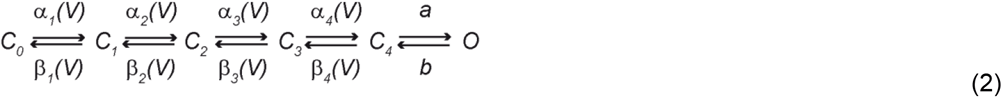

*a* and *b* are the rate constants for transitions between *C*_*4*_ and *O*, and *α*_*i*_*(V)* and *β*_*i*_*(V)* are the voltage dependent forward and backward transition rates that are given by:

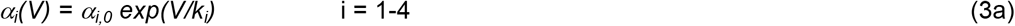

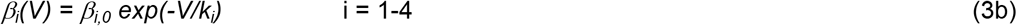

where *α*_*i,0*_ and *β*_*i,0*_ are the forward and backward rate constants at 0 mV and *k*_*i*_ a slope factor.

Ca^2+^ binding to all buffers *(B)* was simulated by second order kinetics with forward and backward binding rate constants *k*_*on*_ and *k*_*off*_, respectively:

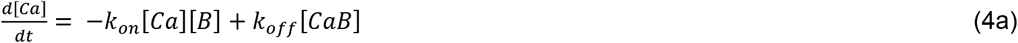

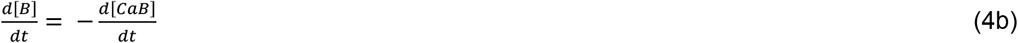

Diffusion of all species (X) was simulated according to

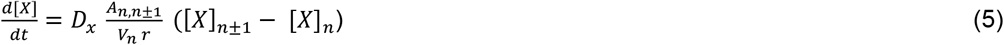

where *D*_*X*_ is the diffusion coefficient, *A* the surface, *V* the volume and *r* the radius of a shell (*n*). Ca^2+^ was cleared by a linear, surface-based extrusion mechanism driven by the difference between [Ca^2+^]_i_ and the resting [Ca^2+^]_i_ (30 nM according to the imaging data).

For fitting the model to the average experimental Ca^2+^ transient, the number of VGCCs was adjusted, yielding 33 VGCCs (**Figure 5**). The other free parameter of the simulation was the extrusion rate, which however affected only the later phase of the decay of the transient. During fitting, the model included the Ca^2+^ indicator dye. Fitting was first performed in a single compartment model (Helmchen and Tank, 2005). Subsequently, fitting was performed in the spatially resolved model with the numbers of VGCCs remaining unaltered and [Ca^2+^]_i_ as reported by the dye was calculated from the concentration of the Ca^2+^-dye complex assuming equilibrium conditions (Schmidt et al., 2003). Since L5PNs are characterized by an immobile and small κ_E_ with unknown binding kinetics, a slow (EB_s_) and a fast immobile endogenous buffer (EB_f_) were assumed (Helmchen et al., 1996; Ohana and Sakmann, 1998; Tran and Stricker, 2018). The fractions of EB_s_ and EB_f_ were adjusted to optimize the fit to the measured Ca^2+^ transient and to sum up to κ_E_ of 25. Following fitting, the indicator dye was removed from the model and the recently developed model for release from L5PN terminals was included (Bornschein et al., 2025). Either the “Syt1 priming model” (Equation 6; **Figure 5**) or the “Syt1 model” (Equation 7; *SI Appendix*, **Figure S5**) were placed at varying distances from the Ca^2+^ sources.

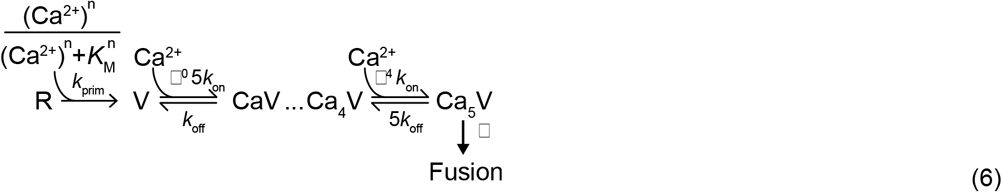

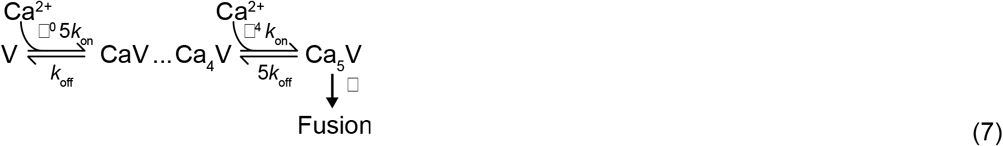

In the priming model (Equation 6), a single Ca^2+^-dependent priming step with Michaelis-Menten kinetics (*K*_M_) and a priming rate *k*_prim_ is assumed. V is the vesicular release sensor of a primed vesicle (Equation 6, 7) that binds a maximum of 5 Ca^2+^ ions. *k*_on_ and *k*_off_ are the forward and backward Ca^2+^ binding rate constants, β^i^ (i=0-4) is a cooperativity factor, and γ the fusion rate. *p*_N_ was obtained by integrating the release rates over time.

A ring like arrangement of the VGCCs around a vesicle was assumed (**Figure 5*D***; ring 1 model in ref. (Bornschein et al., 2019b)). The number of VGCCs driving release was continuously reduced from the maximal value of 33 (**Figure 5*A***) and the coupling distances at which the simulated *p*_N_ values matched the experimentally determined *p*_N_ were obtained. Subsequently, EGTA was included and the consistency between the experimentally quantified EGTA effect and the simulated EGTA effect at a given coupling distance was determined (**Figure 5*B***). The best match between experiments and simulation was obtained for 9 VGCCs at a coupling distance of 51 nm (**Figure 5*C***).

### Quantification and statistical analysis

Data from L5PN-L5PN pairs in PFC were compared to previously published data from these pairs in S1 (**Table 1**)(Bornschein et al., 2019a; Bornschein et al., 2019b). Synaptic responses were determined by fitting a product of two exponential functions to the baseline-subtracted currents, which allows for independent adjustment of the time constants of the rising and falling phases and minimizes noise effects (Bornschein et al., 2013). Synaptic delays were determined from the onset of stimulation to the fitted onset of the EPSC. PPRs were calculated by dividing the second amplitude of two consecutive EPSCs by the first.

Electrophysiological data and Ca^2+^ imaging data were analyzed using custom written routines in Igor Pro (version 6.32 and 8.03, Wavemetrics). Data are shown as median and interquartile range (IQR) or as mean ± standard error (SEM). Summarized data are either shown in boxplots as median ± IQR with mean values included as dashed lines in addition or in bar graphs as mean ± SEM. Normality was tested using the Shapiro-Wilk test. Normally distributed data were compared with the t-test (two groups) or a one-way ANOVA (more than two groups). Non-normally distributed or small samples of data were compared with the Mann-Whitney-U rank sum test (MWU; two groups) or a Kruskal-Wallis ANOVA on ranks (more than two groups). For multiple comparisons post-hoc testing was performed with the Holm-Sidak (one-way ANOVA) or Dunn’s method (ANOVA on ranks). To compare pre- and post-treatment data the paired t-test or the Wilcoxon signed rank test (WSR) was used, depending on the distribution of the data. All statistical tests were two-tailed. P values are indicated as *P<0.05, **P< 0.01 and ***P<0.001. The number of experiments n represents the number of cell pairs, cells or boutons and was chosen sufficiently high to permit reliable statistical analysis. All of the statistical details of experiments can be found in the figures and corresponding figure legends. Statistics were performed with Sigma Plot 11.0 (Dundas Software) and Mathematica 14.

## Acknowledgements

We thank Gudrun Bethge for technical assistance.

This work was supported by a German Research Foundation Grant (SCHM1838/6-1) to H.S..

## Additional information

## Author contributions

Conceptualization: H.S., G.B.; Funding acquisition: H.S.; Methodology: all authors; Investigation: G.B., M.S., A.B., A.A., S.B., H.S.; Data Analysis: G.B., M.S., A.B., A.A., S.B., H.S.; Visualization: G.B., M.S., A.B.; Resources: H.S.; Modeling: H.S.; Supervision: H.S.; Writing – original draft: H.S., G.B.; Writing – review & editing: all authors.

## Data and code availability

All data are available in the main text or in the supplemental information and will be shared by the corresponding author upon request.

All original code has been deposited at Github and is publicly available as of the date of publication.

## Additional Files

### Supplemental information

Document S1. Figures S1–S4, Table S1 and Supplemental references.

## References

Adler EM, Augustine GJ, Duffy SN, Charlton MP (1991) Alien intracellular calcium chelators attenuate neurotransmitter release at the squid giant synapse. J Neurosci 11:1496–1507.

Baur D, Bornschein G, Althof D, Watanabe M, Kulik A, Eilers J, Schmidt H (2015) Developmental tightening of cerebellar cortical synaptic influx-release coupling. J Neurosci 35:1858–1871.

Blatow M, Rozov A, Katona I, Hormuzdi SG, Meyer AH, Whittington MA, Caputi A, Monyer H (2003) A novel network of multipolar bursting interneurons generates theta frequency oscillations in neocortex. Neuron 38:805–817.

Bollmann JH, Sakmann B, Borst JG (2000) Calcium sensitivity of glutamate release in a calyx-type terminal. Science 289:953–957.

Bornschein G, Schmidt H (2019) Synaptotagmin Ca2+ sensors and their spatial coupling to presynaptic Cav channels in central cortical synapses. Front Mol Neurosci 11:494.

Bornschein G, Brachtendorf S, Schmidt H (2019a) Developmental increase of neocortical presynaptic efficacy via maturation of vesicle replenishment. Front Synaptic Neurosci 11:36.

Bornschein G, Eilers J, Schmidt H (2019b) Neocortical high probability release sites are formed by distinct Ca2+ channel-to-release sensor topographies during development. Cell Rep 28:1410–1418 e1414.

Bornschein G, Arendt O, Hallermann S, Brachtendorf S, Eilers J, Schmidt H (2013) Paired-pulse facilitation at recurrent Purkinje neuron synapses is independent of calbindin and parvalbumin during high-frequency activation. J Physiol 591:3355–3370.

Bornschein G, Brachtendorf S, Reinert A, Eshra A, Kraft R, Hirrlinger J, Eilers J, Hallermann S, Schmidt H (2025) The intracellular Ca2+ sensitivity of transmitter release in glutamatergic neocortical boutons. Science 389:48–52.

Borst JG, Helmchen F, Sakmann B (1995) Pre-and postsynaptic whole-cell recordings in the medial nucleus of the trapezoid body of the rat. J Physiol 489:825–840.

Brachtendorf S, Bornschein G, Schmidt H (2025) Estimates of quantal synaptic parameters in light of more complex vesicle pool models. Front Cell Neurosci 19:1556360.

Brandt A, Khimich D, Moser T (2005) Few CaV1.3 channels regulate the exocytosis of a synaptic vesicle at the hair cell ribbon synapse. J Neurosci 25:11577–11585.

Bucurenciu I, Kulik A, Schwaller B, Frotscher M, Jonas P (2008) Nanodomain coupling between Ca2+ channels and Ca2+ sensors promotes fast and efficient transmitter release at a cortical GABAergic synapse. Neuron 57:536–545.

Bullmann T, Kaas T, Ritzau-Jost A, Wohner A, Kirmann T, Rizalar FS, Holzer M, Nerlich J, Puchkov D, Geis C, Eilers J, Kittel RJ, Arendt T, Haucke V, Hallermann S (2024) Human iPSC-Derived Neurons with Reliable Synapses and Large Presynaptic Action Potentials. J Neurosci 44.

Castillo PE (2012) Presynaptic LTP and LTD of excitatory and inhibitory synapses. Cold Spring Harbor perspectives in biology 4:a005728.

Chanda S, Xu-Friedman MA (2010) A low-affinity antagonist reveals saturation and desensitization in mature synapses in the auditory brain stem. J Neurophysiol 103:1915–1926.

Chen JJ, Kaufmann WA, Chen C, Arai I, Kim O, Shigemoto R, Jonas P (2024) Developmental transformation of Ca2+ channel-vesicle nanotopography at a central GABAergic synapse. Neuron.

Clements JD, Silver RA (2000) Unveiling synaptic plasticity: a new graphical and analytical approach. Trends Neurosci 23:105–113.

Douglas RJ, Martin KAC (2004) Neuronal corcuits of the neocortex. Annual Review of Neuroscience 27:419–451.

Doussau F, Schmidt H, Dorgans K, Valera AM, Poulain B, Isope P (2017) Frequency-dependent mobilization of heterogeneous pools of synaptic vesicles shapes presynaptic plasticity. Elife 6:e28935.

Eggermann E, Bucurenciu I, Goswami SP, Jonas P (2012) Nanodomain coupling between Ca2+ channels and sensors of exocytosis at fast mammalian synapses. Nat Rev Neurosci 13:7–21.

Fedchyshyn MJ, Wang LY (2005) Developmental transformation of the release modality at the calyx of Held synapse. J Neurosci 25:4131–4140.

Feldman DE (2009) Synaptic mechanisms for plasticity in neocortex. Annu Rev Neurosci 32:33–55.

Feldman DE (2012) The spike-timing dependence of plasticity. Neuron 75:556–571.

Feldmeyer D (2012) Excitatory neuronal connectivity in the barrel cortex. Frontiers in neuroanatomy 6:24.

Feldmeyer D, Radnikow G (2009) Developmental alterations in the functional properties of excitatory neocortical synapses. J Physiol 587:1889–1896.

Felmy F, Neher E, Schneggenburger R (2003) Probing the intracellular calcium sensitivity of transmitter release during synaptic facilitation. Neuron 37:801–811.

Frick A, Feldmeyer D, Helmstaedter M, Sakmann B (2008) Monosynaptic connections between pairs of L5A pyramidal neurons in columns of juvenile rat somatosensory cortex. Cerebral cortex 18:397–406.

Harlow ML, Ress D, Stoschek A, Marshall RM, McMahan UJ (2001) The architecture of active zone material at the frog’s neuromuscular junction. Nature 409:479–484.

Harris KD, Shepherd GM (2015) The neocortical circuit: themes and variations. Nat Neurosci 18:170–181.

Hefft S, Kraushaar U, Geiger JR, Jonas P (2002) Presynaptic short-term depression is maintained during regulation of transmitter release at a GABAergic synapse in rat hippocampus. J Physiol 539:201–208.

Helmchen F, Tank DW (2005) A single-compartment model of calcium dynamics in nerve terminals and dendrites. In: Imaging neurons: a laboratory manual (Yuste R, Konnerth A, eds), pp 265–275. New York: Cold Spring Harbor Laboratory Press.

Helmchen F, Imoto K, Sakmann B (1996) Ca2+ buffering and action potential-evoked Ca2+ signaling in dendrites of pyramidal neurones. Biophys J 70:1069–1081.

Iwasaki S, Takahashi T (2001) Developmental regulation of transmitter release at the calyx of Held in rat auditory brainstem. J Physiol 534:861–871.

Jackman SL, Regehr WG (2017) The mechanisms and functions of synaptic facilitation. Neuron 94:447–464.

Jarsky T, Tian M, Singer JH (2010) Nanodomain Control of Exocytosis Is Responsible for the Signaling Capability of a Retinal Ribbon Synapse. J Neurosci 30:11885–11895.

Keller D, Babai N, Kochubey O, Han Y, Markram H, Schurmann F, Schneggenburger R (2015) An Exclusion Zone for Ca2+ Channels around Docked Vesicles Explains Release Control by Multiple Channels at a CNS Synapse. PLoS Comput Biol 11:e1004253.

Koike-Tani M, Kanda T, Saitoh N, Yamashita T, Takahashi T (2008) Involvement of AMPA receptor desensitization in short-term synaptic depression at the calyx of Held in developing rats. J Physiol 586:2263–2275.

Kroon T, van Hugte E, van Linge L, Mansvelder HD, Meredith RM (2019) Early postnatal development of pyramidal neurons across layers of the mouse medial prefrontal cortex. Sci Rep 9:5037.

Kusch V, Bornschein G, Loreth D, Bank J, Jordan J, Baur D, Watanabe M, Kulik A, Heckmann M, Eilers J, Schmidt H (2018) Munc13-3 is required for the developmental localization of Ca2+ channels to active zones and the nanopositioning of Cav2.1 near release sensors. Cell Rep 22:1965–1973.

Li L, Bischofberger J, Jonas P (2007) Differential gating and recruitment of P/Q-, N-, and R-type Ca2+ channels in hippocampal mossy fiber boutons. J Neurosci 27:13420–13429.

Lin K-H, Taschenberger H, Neher E (2022) A sequential two-step priming scheme reproduces diversity in synaptic strength and short-term plasticity. Proceedings of the National Academy of Sciences 119:e2207987119.

Markram H, Lübke J, Frotscher M, Roth A, Sakmann B (1997) Physiology and anatomy of synaptic connections between thick tufted pyramidal neurones in the developing rat neocortex. J Physiol 500:409–440.

Matthews G, Fuchs P (2010) The diverse roles of ribbon synapses in sensory neurotransmission. Nat Rev Neurosci 11:812–822.

Matveev V, Zucker RS, Sherman A (2004) Facilitation through buffer saturation: constraints on endogenous buffering properties. Biophys J 86:2691–2709.

Meinrenken CJ, Borst JG, Sakmann B (2002) Calcium secretion coupling at calyx of held governed by nonuniform channel-vesicle topography. J Neurosci 22:1648–1667.

Meyer AC, Neher E, Schneggenburger R (2001) Estimation of quantal size and number of functional active zones at the calyx of held synapse by nonstationary EPSC variance analysis. J Neurosci 21:7889–7900.

Miki T, Malagon G, Pulido C, Llano I, Neher E, Marty A (2016) Actin-and myosin-dependent vesicle loading of presynaptic docking sites prior to exocytosis. Neuron 91:808–823.

Moser T, Beutner D (2000) Kinetics of exocytosis and endocytosis at the cochlear inner hair cell afferent synapse of the mouse. Proc Natl Acad Sci U S A 97:883–888.

Nakamura Y, Harada H, Kamasawa N, Matsui K, Rothman Jason S, Shigemoto R, Silver RA, DiGregorio David A, Takahashi T (2015) Nanoscale distribution of presynaptic Ca2+ channels and its impact on vesicular release during development. Neuron 85:145–158.

Narayanan NS, Hyman JM, Seamans J, Rich EL (2025) Computational Properties of the Prefrontal Cortex. The Journal of Neuroscience 45:e1093252025.

Neher E (2023) Interpretation of presynaptic phenotypes of synaptic plasticity in terms of a two-step priming process. Journal of General Physiology 156.

Neher E, Sakaba T (2001) Combining deconvolution and noise analysis for the estimation of transmitter release rates at the calyx of held. J Neurosci 21:444–461.

Neher E, Brose N (2018) Dynamically primed synaptic vesicle states: Key to understand synaptic short-term plasticity. Neuron 100:1283–1291.

Ohana O, Sakmann B (1998) Transmitter release modulation in nerve terminals of rat neocortical pyramidal cells by intracellular calcium buffers. J Physiol 513:135–148.

Rebola N, Reva M, Kirizs T, Szoboszlay M, Lorincz A, Moneron G, Nusser Z, DiGregorio DA (2019) Distinct nanoscale calcium channel and synaptic vesicle topographies contribute to the diversity of synaptic function. Neuron 104:693–710 e699.

Regehr WG, Carey MR, Best AR (2009) Activity-dependent regulation of synapses by retrograde messengers. Neuron 63:154–170.

Reyes A, Sakmann B (1999) Developmental switch in the short-term modification of unitary EPSPs evoked in layer 2/3 and layer 5 pyramidal neurons of rat neocortex. J Neurosci 19:3827–3835.

Rozov A, Burnashev N, Sakmann B, Neher E (2001) Transmitter release modulation by intracellular Ca2+ buffers in facilitating and depressing nerve terminals of pyramidal cells in layer 2/3 of the rat neocortex indicates a target cell-specific difference in presynaptic calcium dynamics. J Physiol 531:807–826.

Sabatini BL, Oertner TG, Svoboda K (2002) The life cycle of Ca2+ ions in dendritic spines. Neuron 33:439–452.

Scheuss V, Schneggenburger R, Neher E (2002) Separation of presynaptic and postsynaptic contributions to depression by covariance analysis of successive EPSCs at the calyx of held synapse. J Neurosci 22:728–739.

Schmidt H (2019) Control of presynaptic parallel fiber efficacy by activity-dependent regulation of the number of occupied release sites. Frontiers in systems neuroscience 13:30.

Schmidt H, Stiefel K, Racay P, Schwaller B, Eilers J (2003) Mutational analysis of dendritic Ca2+ kinetics in rodent Purkinje cells: role of parvalbumin and calbindin D28k. J Physiol (Lond) 551:13–32.

Schmidt H, Brachtendorf S, Arendt O, Hallermann S, Ishiyama S, Bornschein G, Gall D, Schiffmann SN, Heckmann M, Eilers J (2013) Nanodomain coupling at an excitatory cortical synapse. Curr Biol 23:244–249.

Schneggenburger R, Neher E (2000) Intracellular calcium dependence of transmitter release rates at a fast central synapse. Nature 406:889–893.

Scimemi A, Diamond JS (2012) The number and organization of Ca2+ channels in the active zone shapes neurotransmitter release from Schaffer collateral synapses. J Neurosci 32:18157–18176.

Shahrezaei V, Cao A, Delaney KR (2006) Ca2+ from one or two channels controls fusion of a single vesicle at the frog neuromuscular junction. J Neurosci 26:13240–13249.

Silver RA (2003) Estimation of nonuniform quantal parameters with multiple-probability fluctuation analysis: theory, application and limitations. J Neurosci Methods 130:127–141.

Singer JH, Diamond JS (2003) Sustained Ca2+ entry elicits transient postsynaptic currents at a retinal ribbon synapse. J Neurosci 23:10923–10933.

Taschenberger H, Leao RM, Rowland KC, Spirou GA, von Gersdorff H (2002) Optimizing synaptic architecture and efficiency for high-frequency transmission. Neuron 36:1127–1143.

Tran V, Stricker C (2018) Diffusion of Ca2+ from small boutons en passant into the axon shapes AP-evoked Ca2+ transients. Biophys J 115:1344–1356.

Vyleta NP, Jonas P (2014) Loose coupling between Ca2+ channels and release sensors at a plastic hippocampal synapse. Science 343:665–670.

Weichard I, Taschenberger H, Gsell F, Bornschein G, Ritzau-Jost A, Schmidt H, Kittel RJ, Eilers J, Neher E, Hallermann S, Nerlich J (2023) Fully-primed slowly-recovering vesicles mediate presynaptic LTP at neocortical neurons. Proc Natl Acad Sci U S A 120:e2305460120.

Yarim A, Brachtendorf B, Schmidt H, Bornschein G (2026) Pyramidal neuron synapses in M2 exhibit properties intermediate between prefrontal cortex and M1 synapses. in preparation

Zucker RS, Regehr WG (2002) Short-term synaptic plasticity. Annual review of physiology 64:355–405.

