## Supplemental Information for "Loose coupling between Ca^2+^ channels and release sensors as a synaptic correlate of higher order brain function"

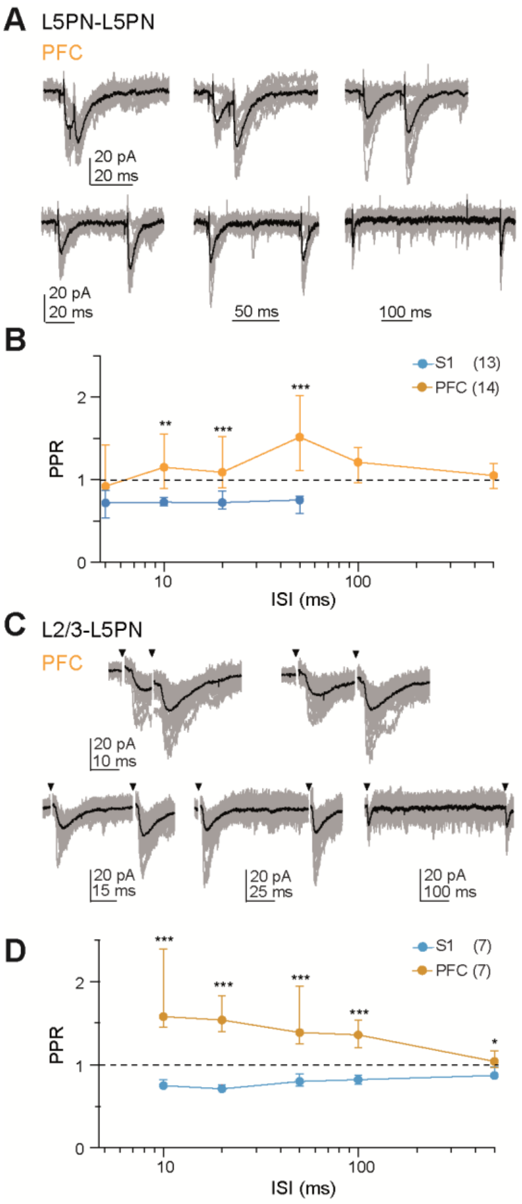


**Figure S1. Short-term plasticity in PFC and S1.**

**(*A*)** EPSCs recorded from a L5PN-L5PN pair in PFC at interstimulus intervals (ISIs) ranging from 5 to 500 ms.

**(*B*)** Comparison of PPRs recorded at different ISIs from pairs of L5PNs in PFC (orange) and S1 (blue). Averages shown as medians and IQRs (n in brackets; P=0.104, **P=0.008, ***P<0.001, MWU). S1 data are from (Bornschein et al., 2019).

**(*C*)** Same as in (***A***), but for recordings from prefrontal L5PNs after extracellular stimulation in L2/3 with paired-pulses at ISIs of 10 to 500 ms.

**(*D*)** Same as in (***B***), but for L2/3-L5PN synapses in PFC and S1 (***P<0.001, MWU).

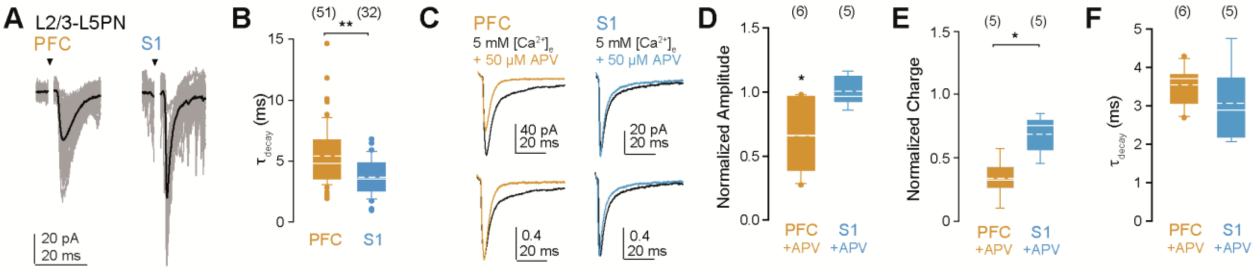


**Figure S2. Different NMDA receptor contributions in PFC and S1.**

(***A***) Examples of EPSCs recorded from L2/3-L5PN connections in PFC and S1.
(***B***) Summary of EPSC decay time constants ($\tau_{decay}$) in PFC and S1 (**P=0.002, MWU).
(***C***) Top: Averaged example recordings before (baseline in black) and after application of 50 μM APV in PFC (orange) and S1 (blue). Bottom: EPSC recordings from top graphs normalized to baseline emphasizing changes in synaptic decay.
(***D-F***) Box plots comparing normalized EPSC amplitudes (***D***; *P=0.031, P=1.0, WSR), their charges (***E***; *P=0.016, MWU) and $\tau_{decay}$ values in the presence of 50 μM APV (***F***; P=0.329) in PFC and S1.

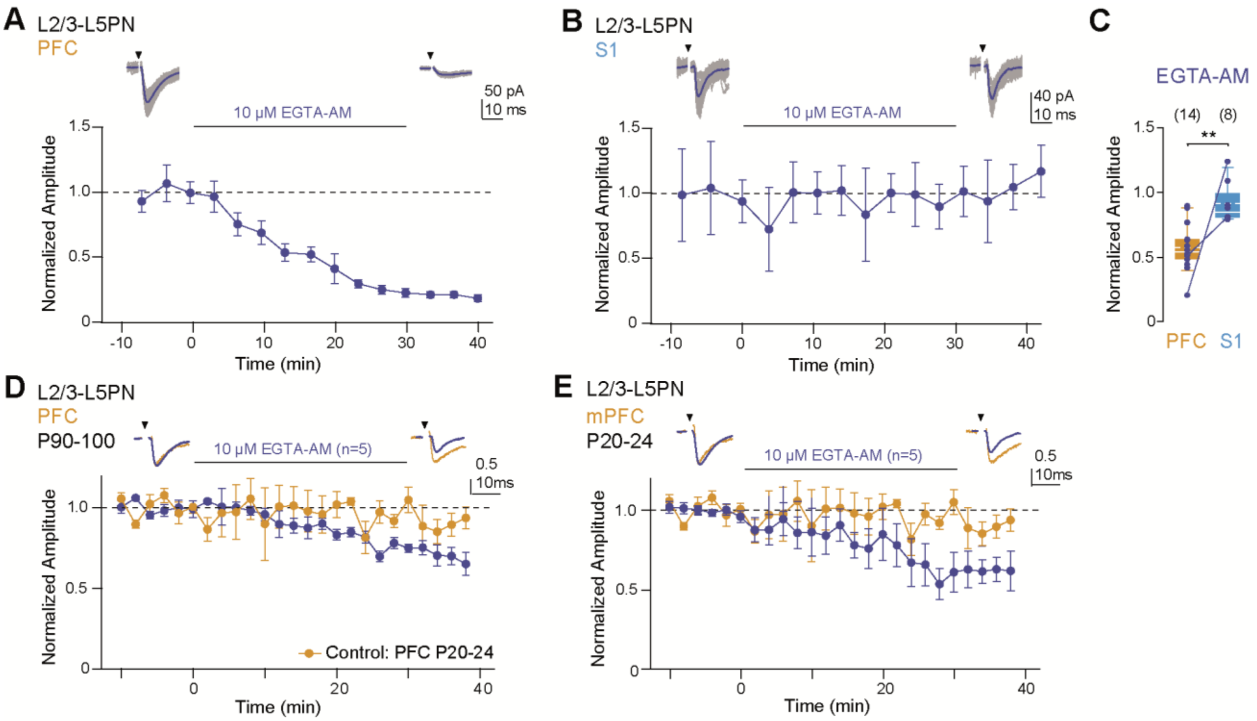


**Figure S3. EGTA sensitivity in PFC.**

(***A, B***) Example recording of baseline-normalized EPSC amplitudes (means ± SDs, 20 EPSCs each were binned) recorded from a L2/3-L5PN connection in PFC (**A**) and S1 cortex (**B**) derived from the same mouse (P24). Following baseline recordings (10 min), slices were perfused for 30 min (solid line) with ACSF containing 10 µM EGTA-AM (blue), and thereafter rinsed for 10 min with ACSF (test period). Insets: EPSCs (averages in blue, individual recordings in gray) during baseline and test period.
(***C***) Box plot summarizing EGTA-AM effects in PFC and S1. Individual experiments are included as data points, data obtained from the same mouse were connected by a line. Note, the significant EGTA sensitivity in PFC (**P=0.002, MWU).
(***D***) Averaged, baseline-normalized EPSCs (2 min bins) recorded from L2/3-L5PN connections in PFC of 90-100 days old mice as in **Figure 2*C***. Insets: Averaged EPSCs during baseline and test period.
(***E***) Same as in (***D***), but for L2/3-L5PN connections in mPFC of 21-26 days old mice (**Figure 2*D***).

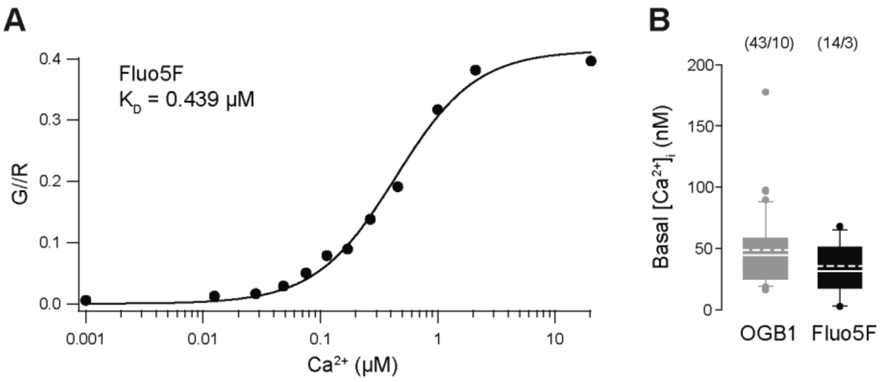


**Figure S4. *In vitro* calibration curve and basal $\Delta[Ca^{2+}]_i$.**

**(*A*)** Calibration curve of Fluo-5F in a K-gluconat-based pipette solution. The $K_D$ value of 439 nM was derived from a Hill fit (solid curve).

**(*B*)** Comparison of basal $[Ca^{2+}]_i$ determined with Fluo-5F and OGB-1, respectively (P=0.204, MWU)

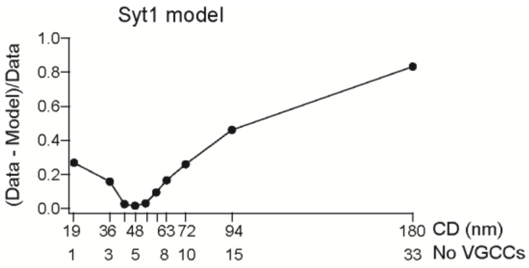


**Figure S5. Prediction of the coupling distance with an alternative Syt1 model.**

As in **Figure 5*B*** but instead of the Syt1 priming model, the Syt1 model (Bornschein et al., 2019; Bornschein et al., 2025) was used for simulating the release rates and the corresponding $p_N$ values.

**Table S1. Model Parameters.**

| Parameters | | Values | References/Notes |
|---|---|---|---|
| Bouton volume | | $0.13 * 10^{-15}$ l | (Schikorski and Stevens, 1999)[1] |
| Calcium | $[Ca^{2+}]_{rest}$ | $30*10^{-9}$ M | Measured |
| | $D$ | $220 *10^{-12}$ $m^2s^{-1}$ | (Allbritton et al., 1992) |
| Fluo-5F | [Fluo] | $140 *10^{-6}$ M | 70% of pipette concentration |
| | $K_D$ | $439 *10^{-9}$ M | Measured |
| | $k_{off}$ | 300 s-1 | (Scott and Rusakov, 2006) |
| | $D$ | $220 *10^{-12}$ $m^2s^{-1}$ | [2] |
| EGTA | $K_D$ | $7*10^{-8}$ M | (Nägerl et al., 2000) |
| | $k_{off}$ | $0.7\ s^{-1}$ | ibid. |
| | $k_{on}$ | $1*10^7\ M^{-1}s^{-1}$ | ibid. |
| | $D$ | $220 *10^{-12}$ $m^2s^{-1}$ | (Naraghi and Neher, 1997) |
| ATP | [ATP] | $6 *10^{-4}$ M | Pipette concentration |
| | $K_D$ | $2*10^{-4}$ M | (Meinrenken et al., 2002) |
| | $k_{off}$ | $100\ 000\ s^{-1}$ | ibid. |
| | $k_{on}$ | $5*10^8\ M^{-1}s^{-1}$ | ibid. |
| | $D$ | $220 *10^{-12}$ $m^2s^{-1}$ | ibid. |
| | $\kappa$ | 3 | Calculated |
| Immobile endogenous $Ca^{2+}$ buffer (EB) | $\kappa_E$ | 25 | (Tran and Stricker, 2018; Bornschein et al., 2019) |
| Fast EB | $[EB_f]$ | $46.5 *10^{-6}$ M | Calculated |
| | $K_D$ | $2*10^{-6}$ M | (Vyleta and Jonas, 2014; Bornschein et al., 2019) |
| | $k_{off}$ | $1000\ s^{-1}$ | ibid. |
| | $k_{on}$ | $5*10^8\ M^{-1}s^{-1}$ | ibid. |
| | $\kappa_E$ | 23.25 | Calculated |
| Slow EB | $[EB_s]$ | $1.23*10^{-7}$ M | Calculated |
| | $K_D$ | $7*10^{-8}$ M | (Nägerl et al., 2000) |
| | $k_{off}$ | $0.7\ s^{-1}$ | ibid. |
| | $k_{on}$ | $1*10^7\ M^{-1}s^{-1}$ | ibid. |
| | $\kappa_E$ | 1.75 | Calculated |
| P/Q-type channel | a, b | 247.71, 8.28 $ms^{-1}$ | (Li et al., 2007) |
| | $\alpha_{1,0}$, $\beta_{1,0}$ | 5.89, 14.99 $ms^{-1}$ | ibid. |
| | $\alpha_{2,0}$, $\beta_{2,0}$ | 9.21, 6.63 $ms^{-1}$ | ibid. |
| | $\alpha_{3,0}$, $\beta_{3,0}$ | 5.2, 132.8 $ms^{-1}$ | ibid. |
| | $\alpha_{4,0}$, $\beta_{4,0}$ | 1823.18, 248.58 $ms^{-1}$ | ibid. |
| | k1 | 62.61 mV | ibid. |
| | k2 | 33.92 mV | ibid. |
| | k3 | 135.08 mV | ibid. |
| | k4 | 20.86 mV | ibid. |
| Release sensor models | | | (Bornschein et al., 2025) |
| Syt1 model | $k_{off}$ | $10\ 000\ s^{-1}$ | ibid. |
| | $k_{on}$ | $1*10^8\ M^{-1}s^{-1}$ | ibid. |
| | $\gamma$ | $5900\ s^{-1}$ | ibid. |

| | | | |
|---|---|---|---|
| | $\beta$ | 4 | ibid. |
| Syt1 priming model | $k_{off}$ | 10 000 $s^{-1}$ | ibid. |
| | $k_{on}$ | $1*10^8$ $M^{-1}s^{-1}$ | ibid. |
| | $\gamma$ | 4000 $s^{-1}$ | ibid. |
| | $\beta$ | 4 | ibid. |
| | $K_M$ | $20*10^{-6}$ M | ibid. |
| | $k_{prim}$ | 6000 $s^{-1}$ | ibid. |
| | n | 5 | ibid. |

[1] Corrected for the space occupied by vesicles and organelles (Wilhelm et al., 2014).
[2] Assumed to be similar to free $Ca^{2+}$ (Bucurenciu et al., 2008).

### Supplemental References


Allbritton NL, Meyer T, Stryer L (1992) Range of messenger action of calcium ion and inositol 1,4,5-trisphosphate. Science 258:1812-1815.

Bornschein G, Eilers J, Schmidt H (2019) Neocortical high probability release sites are formed by distinct $Ca^{2+}$ channel-to-release sensor topographies during development. Cell Rep 28:1410-1418 e1414.

Bornschein G, Brachtendorf S, Reinert A, Eshra A, Kraft R, Hirrlinger J, Eilers J, Hallermann S, Schmidt H (2025) The intracellular $Ca^{2+}$ sensitivity of transmitter release in glutamatergic neocortical boutons. Science 389:48-52.

Bucurenciu I, Kulik A, Schwaller B, Frotscher M, Jonas P (2008) Nanodomain coupling between $Ca^{2+}$ channels and $Ca^{2+}$ sensors promotes fast and efficient transmitter release at a cortical GABAergic synapse. Neuron 57:536-545.

Li L, Bischofberger J, Jonas P (2007) Differential gating and recruitment of P/Q-, N-, and R-type $Ca^{2+}$ channels in hippocampal mossy fiber boutons. J Neurosci 27:13420-13429.

Meinrenken CJ, Borst JG, Sakmann B (2002) Calcium secretion coupling at calyx of held governed by nonuniform channel-vesicle topography. J Neurosci 22:1648-1667.

Nägerl UV, Novo D, Mody I, Vergara JL (2000) Binding kinetics of calbindin-$D_{28k}$ determined by flash photolysis of caged $Ca^{2+}$. Biophys J 79:3009-3018.

Naraghi M, Neher E (1997) Linearized buffered $Ca^{2+}$ diffusion in microdomains and its implications for calculation of $[Ca^{2+}]$ at the mouth of a calcium channel. J Neurosci 17:6961-6973.

Schikorski T, Stevens CF (1999) Quantitative fine-structural analysis of olfactory cortical synapses. Proc Natl Acad Sci U S A 96:4107-4112.

Scott R, Rusakov DA (2006) Main determinants of presynaptic $Ca^{2+}$ dynamics at individual mossy fiber-CA3 pyramidal cell synapses. J Neurosci 26:7071-7081.

Tran V, Stricker C (2018) Diffusion of $Ca^{2+}$ from small boutons en passant into the axon shapes AP-evoked $Ca^{2+}$ transients. Biophys J 115:1344-1356.

Vyleta NP, Jonas P (2014) Loose coupling between $Ca^{2+}$ channels and release sensors at a plastic hippocampal synapse. Science 343:665-670.

Wilhelm BG, Mandad S, Truckenbrodt S, Kröhnert K, Schäfer C, Rammner B, Koo SJ, Claßen GA, Krauss M, Haucke V, Urlaub H, Rizzoli SO (2014) Composition of isolated synaptic boutons reveals the amounts of vesicle trafficking proteins. Science 344:1023-1028.
